# T cells regulate intestinal motility and shape enteric neuronal responses to intestinal microbiota

**DOI:** 10.1101/2024.05.23.595563

**Authors:** Patricia Rodrigues Marques de Souza, Catherine M. Keenan, Laurie E. Wallace, Yasaman Bahojb Habibyan, Marcela Davoli-Ferreira, Christina Ohland, Fernando A. Vicentini, Kathy D. McCoy, Keith A. Sharkey

## Abstract

The gut microbiota and immune system maintain intestinal homeostasis and regulate gut physiology in concert with the enteric nervous system (ENS). However, the underlying mechanisms remain incompletely understood. Using wildtype and T-cell deficient germ-free mice colonized with segmented filamentous bacteria (SFB) or specific pathogen-free (SPF) microbiota, we studied immune regulation of the ENS and intestinal motility. Colonization markedly increased Th17 cells and Treg expressing RORγ^+^T cells in both the ileum and colon of wildtype mice. T cells were necessary for the normalization of intestinal motility after colonization by SPF microbiota, and for SFB to restore neuronal density in the ENS of the ileum of germ-free mice. T cells were also required for neurogenic responses in myenteric neurons of the ileum, but not the colon, and for regulating the levels of nestin expression. The cytokines IL-1β and IL-17A mediate the enteric neurogenic response to an SPF microbiota but were not involved in the regulation of intestinal motility. Together, our findings provide new insights into the microbiota-neuroimmune dialogue that regulates intestinal physiology.

---

The gastrointestinal (GI) tract is regulated by a complex nervous system that lies in the wall of the gut called the enteric nervous system (ENS)^1, 2^. The ENS controls the physiological functions of the gut including GI motility, fluid and electrolyte secretion, immune function, mucosal growth, and intestinal permeability^1, 2^. It consists of neurons and glia arranged in two ganglionated plexuses, the myenteric and submucosal plexuses^1, 2, 3^. A network of nerve fibres and glial processes innervate cellular targets throughout the gut wall including epithelial cells, glands, immune cells, and smooth muscle^1, 2, 3^.

The gut microbiota programs and shapes the development of the ENS and in doing so, is in part responsible for the regulation of gut physiology^4, 5, 6, 7^. In adult mice, depleting the gut microbiota with a cocktail of antibiotics leads to a reduction in the numbers of enteric neurons in the myenteric and submucosal plexuses, and small bowel motility is slowed^8, 9, 10, 11, 12, 13, 14^. The effects of antibiotic treatment are completely reversed by natural recolonization by the gut microbiota^8, 10^. The mechanisms involved in this response remain to be fully elucidated.

Germ-free mice also have slowed intestinal transit and abnormalities in their enteric innervation, but various studies report different results in terms of the density of enteric neurons in different regions of the GI tract^9, 11, 14, 15, 16, 17, 18, 19, 20, 21^. Most studies report a reduced number of neurons compared to conventionally raised (specific-pathogen free [SPF]) mice^9, 11, 14, 15, 20^. Upon recolonization, the number of neurons is generally found to return to control levels^15, 20^. However, Yan *et al.*, found that recolonization reduced the enteric neuronal density in the colon; with greater effects seen in mice monocolonized with a commensal bacterium (*Clostridium ramosum*) known to stimulate RORγ^+^T cells (Th17 cells)^19^. Moreover, they showed that microbial signals from commensal microbiota that induce RORγ^+^ Treg cells condition neuronal density^19^. However, the microbial and immune mechanisms that underlie the enteric neurogenic response in adult animals to bacterial colonization are not fully understood.

A number of recent studies of the ENS have revealed potential mechanisms for enteric neurons to interact with T cells via mediators that are typically regarded as immune signaling molecules^19, 22, 23, 24^. We were intrigued by the observation that single cell sequencing studies revealed that enteric neurons express the IL-17 receptor (IL-17R; *Il17ra-e*)^25, 26^, the prototypic cytokine of Th17 cells. Given the potential for interactions of the ENS with T cells and their importance, we hypothesized that restoration of enteric neuronal homeostasis and normalization of small intestinal transit after microbiota colonization of germ-free mice are dependent on T cells and specific cytokines.

Previous studies have determined that adult neurogenesis occurs through enteric glial dedifferentiation of Sox2 expressing enteric glia, or the stimulation of progenitor cells, which express markers including nestin, doublecortin-like kinase 1 and/or P75 neurotrophin receptor^27, 28, 29, 30, 31, 32, 33, 34, 35^. Here we investigated the expression of Sox2 and nestin in enteric neurons of the mouse ENS, and intestinal motility in three conditions: the germ-free state, monocolonization with the potent inducer of Th17 cells, segmented filamentous bacteria (SFB, *Candidatus savagella*)^36^, and colonization with an SPF microbiota that contains a typical mouse microbiota including SFB. To conduct these studies, we made use of adult wildtype germ-free and germ-free T cell-deficient mice, that were monocolonized with SFB or colonized with an SPF microbiota.

We show that T cells are important for the normalization of intestinal motility after colonization and that the cytokines IL-1β and IL-17A mediate the enteric neurogenic response to an SPF microbiota. SFB monocolonization stimulates enteric neurogenesis, requiring T cells in the small intestine, but not the colon, but it is not sufficient to normalize small intestinal motility. Together, our findings provide new insights into the microbiota-neuroimmune dialogue that regulates intestinal physiology, highlighting a role for T cells in regulation of intestinal motility and the cytokines IL-1β and IL-17A in enteric neurogenesis that occurs in adult mice in response to microbial colonization of the gut.

## Results

### Small Intestinal transit

Since small intestinal transit in mice is measured as a proportion of the length of the gut, we first assessed the length of the small intestine (and the colon [Supplementary Results]) of the mice used in this study. Small intestinal length did not differ between the groups of wildtype mice, ranging from 36.6 ± 0.5cm – 37.2 ± 1.0cm in the 3 groups (n=10/group). The length of the small intestine of the SPF-colonized mice (36.6 ± 0.5cm) was virtually identical to that of conventional mice raised under SPF conditions from birth (38.7 ± 0.7cm [n=10]). As observed with wildtype germ-free mice, small intestinal length did not differ between the groups of T cell-deficient mice (ranging from 38.3 ± 0.7cm – 38.8 ± 0.6cm [n=8/group; two-way ANOVA, P>0.05]). These data reveal that the lengths of the small intestine are comparable in all groups.

We then determined small intestinal transit by the passage of gavaged carmine red dye. Wildtype germ-free mice and germ-free mice monocolonized with SFB had similar transit times (Figure 1). In contrast, wildtype germ-free mice colonized with SPF had a significantly faster small intestinal transit compared to germ-free mice (Figure 1), which was similar to that observed in mice housed from birth in a conventional facility (74.0±3.5% [n=10], dashed line, Figure 1). These data reveal that SFB monocolonization was not sufficient to restore small intestinal transit to normal levels. Colonization of T cell-deficient germ-free mice with either SFB or SPF did not alter their small intestinal transit (ANOVA, P>0.05) (Figure 1). These data show that T cells are required for the normalization of small intestinal transit that occurs when the gut is colonized with an SPF microbiota.

**Figure 1.**
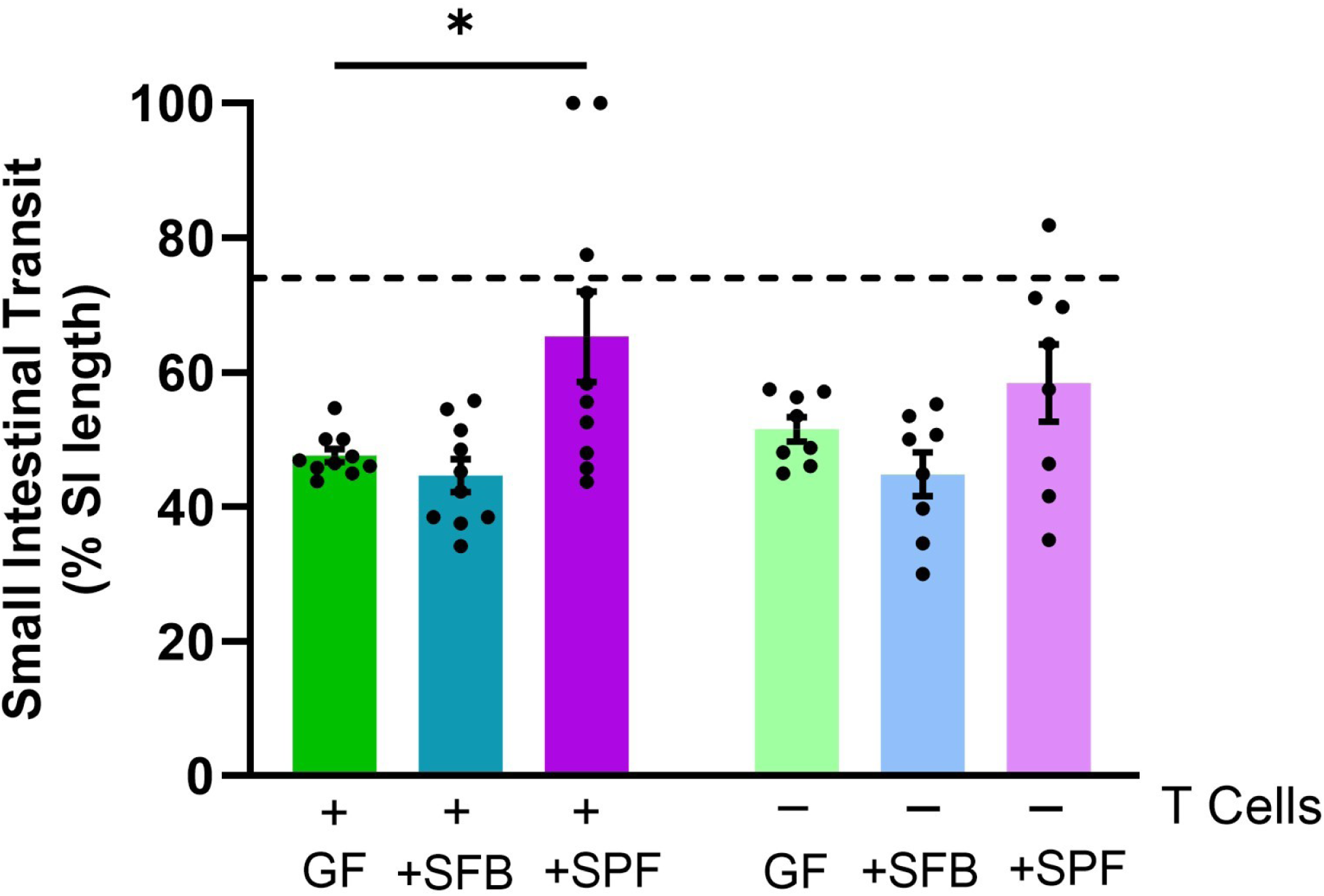
Small intestinal transit is modulated by the gut microbiota in a T cell-dependent manner. Small intestinal transit was measured after oral gavage of carmine red in wildtype germ-free mice (GF), GF mice colonized with SFB or SPF microbiota (+), T cell-deficient GF mice, and T cell-deficient GF mice colonized with SFB or SPF microbiota (-). Each dot represents an individual animal. Dashed line indicates average small intestinal transit measured in wildtype mice colonized with SPF from birth. Data are pooled from 2-4 independent experiments run for each genotype. Data were analyzed using two-way ANOVA, followed by Šídák’s multiple comparison test. N=8-10/group. * p<0.05.

### Enteric neuronal density

Next, we assessed the density of neurons in the submucosal and myenteric plexuses as determined by counts of HuC/D cells in whole-mount preparations of the ileum and colon. In both regions of the gut and in both plexuses, we found that wildtype germ-free mice had significantly reduced numbers of neurons compared to mice colonized with an SPF microbiota (Figure 2). The numbers of neurons in the ENS of SPF-colonized wildtype mice were very similar to that in wildtype mice born with an SPF microbiota (dashed line, Figure 2) and to those we have previously reported^8^ (Figure 2). In wildtype mice colonized with SFB, similar numbers of neurons to those of SPF-colonized mice are observed in both plexuses and both regions of the gut, despite SFB primarily colonizing the ileum^36, 37^. Thus, both SFB and SPF microbiota are sufficient to stimulate enteric neurogenesis and the restoration of neuronal density in adult wildtype germ-free mice.

**Figure 2.**
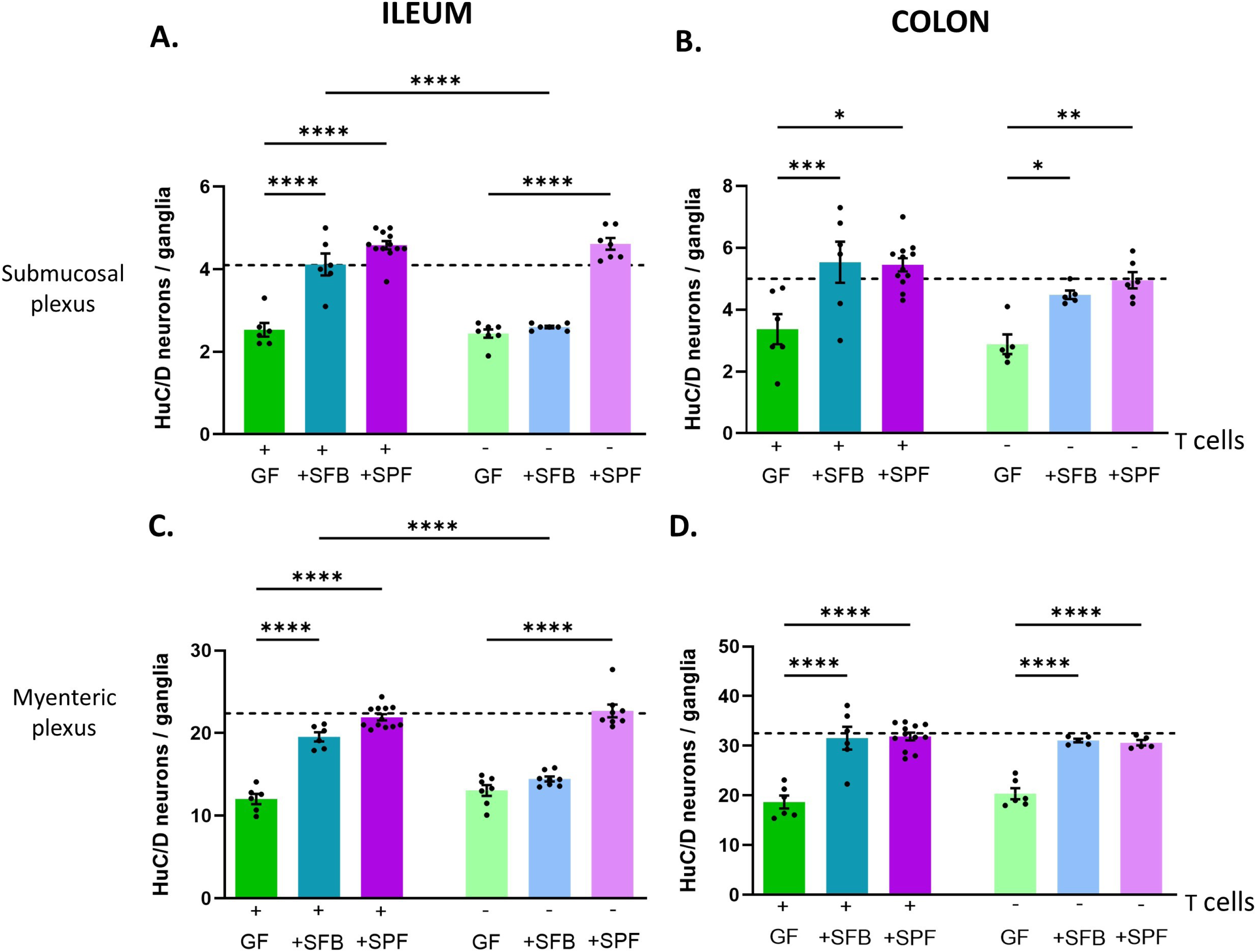
Enteric neuronal density is modulated by the microbiota in a T cell– and region-specific manner. Enteric neuronal density was assessed using counts of HuC/D neurons in the submucosal and myenteric plexuses in wildtype germ-free mice (GF), GF mice colonized with SFB or SPF microbiota (+), T cell-deficient GF mice, and T cell-deficient GF mice colonized with SFB or SPF microbiota (-) in the ileum (**A, C**) and colon (**B, D**). Each dot represents an individual animal. Dashed line indicates average neuronal density measured in wildtype mice colonized with SPF from birth. Data are pooled from 2-4 independent experiments run for each genotype. Data were analyzed using two-way ANOVA, followed by Šídák’s multiple comparison test. N=4-12/group. * p<0.05, ** p<0.01, *** p<0.001, **** p<0.0001.

The neuronal density in the submucosal and myenteric plexuses of the ileum and colon of T cell-deficient germ-free mice were similar to those observed in wildtype germ-free mice (Figure 2). Colonization with SPF microbiota in T cell-deficient mice significantly increased the neuronal density in the submucosal and myenteric plexuses of both the ileum and colon to similar levels as in the wildtype animals (Figure 2). In contrast, when T cell-deficient germ-free mice were monocolonized with SFB, we observed region-specific differences in the responses (Figure 2). In both the submucosal and myenteric plexuses of the ileum, but not the colon, SFB colonization of T cell-deficient mice failed to restore neuronal density, suggesting that T cells are required for the neurogenic response to SFB in the small intestine, but not in the colon.

### Enteric neurogenesis

To further investigate enteric neurogenesis, we examined the presence of Sox2 and nestin in enteric neurons, since they are established markers of adult neurogenesis in the myenteric plexus^27, 28, 30, 34^. In the submucosal plexuses there is little Sox2 expression (Supplementary Figure 1) and no nestin expression, suggesting that other markers are needed to determine the source of neurons in the submucosal plexus, and we did not investigate this further.

In the myenteric plexus of wildtype and T cell-deficient germ-free mice, there were very few Sox2-HuC/D cells in the ileum, but in the colon, 1-2 cells per ganglion were observed in both groups of mice (Figure 3). In the myenteric plexus of the ileum of wildtype mice, SPF colonization significantly increased the number of Sox2-HuC/D cells, a response that was completely absent in T cell-deficient mice (P<0.001, two-way ANOVA, Figure 3, upper panel). In contrast, in the colonic myenteric plexus, there was a significant increase in Sox2-HuC/D cells in both wildtype and T cell-deficient mice in response to SPF colonization. In the myenteric plexus of the ileum of wildtype mice, SFB colonization significantly increased the number of Sox2-HuC/D cells, but no Sox2-HuC/D cells were observed in T cell-deficient mice (Figure 3). In contrast, in the colonic myenteric plexus, there was a significant increase in Sox2-HuC/D cells in both wildtype and T cell-deficient mice monocolonized with SFB (Figure 3, lower panel), illustrating important regional differences.

**Figure 3.**
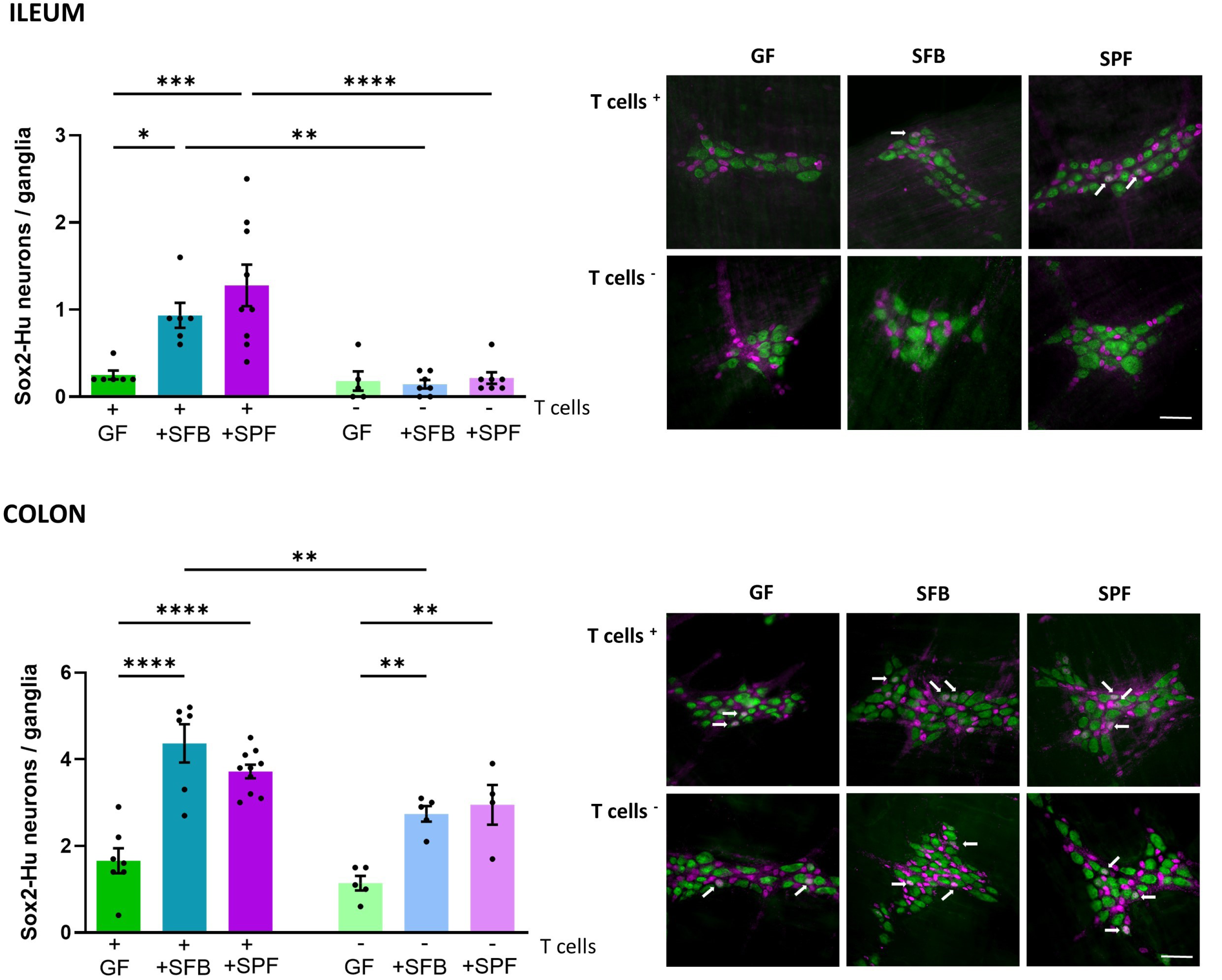
Sox2 in myenteric neurons is modulated by the microbiota in a T cell– and region-specific manner. Left panels. Sox2-HuC/D neurons in the myenteric plexus of wildtype germ-free mice (GF), GF mice colonized with SFB or SPF microbiota (+), T cell-deficient GF mice, and T cell-deficient GF mice colonized with SFB or SPF microbiota (-) in the ileum (upper panels) and colon (lower panels). Each dot represents an individual animal. Data are pooled from 2-4 independent experiments run for each genotype. Data were analyzed using two-way ANOVA, followed by Šídák’s multiple comparison test. N=4-10/group. * p<0.05, ** p<0.01, *** p<0.001, **** p<0.0001. Right panels. Immunohistochemical double-labeling of Sox2 (purple) and HuC/D (green) in the ileum and colon of GF, SFB, and SPF colonized wildtype (+) and T cell-deficient (-) mice. Examples of doubled-labelled neurons are shown using arrows. Scale bar: 30μm.

In the myenteric plexus of the ileum of wildtype mice, nestin-HuC/D cells significantly increased after both SFB and SPF colonization (Figure 4). Curiously, the ileum of T cell-deficient mice contained higher numbers of nestin^+^ cells in all groups of mice and no evidence that colonization had any effect on the numbers of nestin-HuC/D cells (Figure 4). There was a significant increase in nestin expression in the colonic myenteric plexus of wildtype mice colonized with SFB (Figure 4, lower panel). As in the ileum, in the colonic myenteric plexus of T cell-deficient mice, the baseline level of nestin expression was increased and there was no evidence that colonization altered the numbers of nestin-HuC/D cells (Figure 4, lower panel).

**Figure 4.**
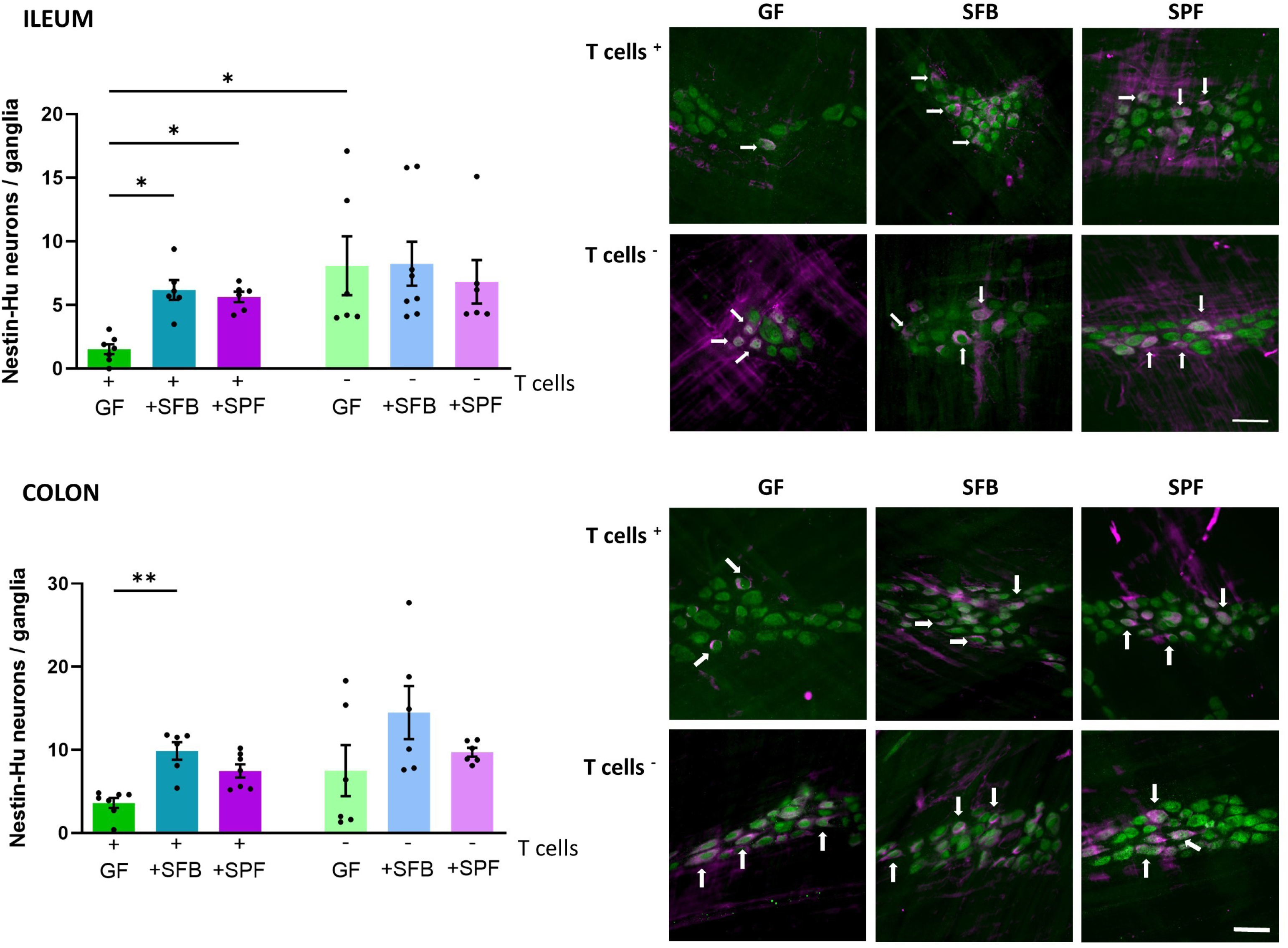
Nestin in myenteric neurons is modulated by the microbiota in wildtype, but not T cell-deficient mice. Left panels. Nestin-HuC/D neurons in the myenteric plexus of wildtype germ-free mice (GF), GF mice colonized with SFB or SPF microbiota (+), T cell-deficient GF mice, and T cell-deficient GF mice colonized with SFB or SPF microbiota (-) in the ileum (upper panels) and colon (lower panels). Each dot represents an individual animal. Data are pooled from 2-4 independent experiments run for each genotype. Data were analyzed using two-way ANOVA, followed by Šídák’s multiple comparison test. N=5-8/group. * p<0.05, ** p<0.01. Right panels. Immunohistochemical double-labeling of nestin (purple) and HuC/D (green) in the ileum and colon of GF, SFB, and SPF colonized wildtype (+) and T cell-deficient (-) mice. Examples of doubled-labelled neurons are shown using arrows. Scale bar: 30μm.

Together, these data reveal a region-specific neuronal regulation of the ENS by the gut microbiota, involving both de-differentiation of enteric glia and stimulation of enteric progenitor cells, based on the presence of Sox2 and nestin in neurons, respectively. Interestingly, T cells are required for regulating Sox2 expression in the ileum, but not the colon, and for regulating the levels of nestin expression in enteric neurons of the myenteric plexus.

### Microbiota profiling after colonization with SPF microbiota and SFB

We next wanted to ascertain whether mice that had been colonized with an SPF microbiota in the presence or absence of T cells had altered microbial composition in the gut, by examining the cecal microbiota. We measured alpha diversity using the Simpson, Chao1 and Shannon indices and found no significant differences between SPF-colonized germ-free and SPF-colonized germ-free T cell-deficient mice (Figure 5A). Beta diversity analysis using the Bray-Curtis distance metric showed that samples clustered separately and differed significantly when analyzed using PERMANOVA (Figure 5B). To visualize microbial composition, we generated the top 30 prevalent amplicon sequencing variants (ASVs) to show relative abundance (Supplementary Figure 2). From here, we sought to better visualize differences in ASVs between the SPF-colonized germ-free and SPF-colonized germ-free T cell-deficient mice, so we examined differential abundance between groups using DeSeq2 (Supplementary Figure 3). Following colonization, T cell-deficient mice showed a significant decrease in two ASVs, Roseburia and Dubosiella, both belonging to the Firmicutes phylum, when compared to wildtype mice. Furthermore, in T cell-deficient mice 12 significantly increased ASVs, most belonging to the Firmicutes phylum, followed by the Bacteriodeta phylum were observed (Supplementary Figure 3). Expectedly, Firmicutes and Bacteriodeta are the two most prevalent phyla within the gastrointestinal tract^38^. These data show that overall, the cecal microbiota was significantly altered in the absence of T cells.

**Figure 5.**
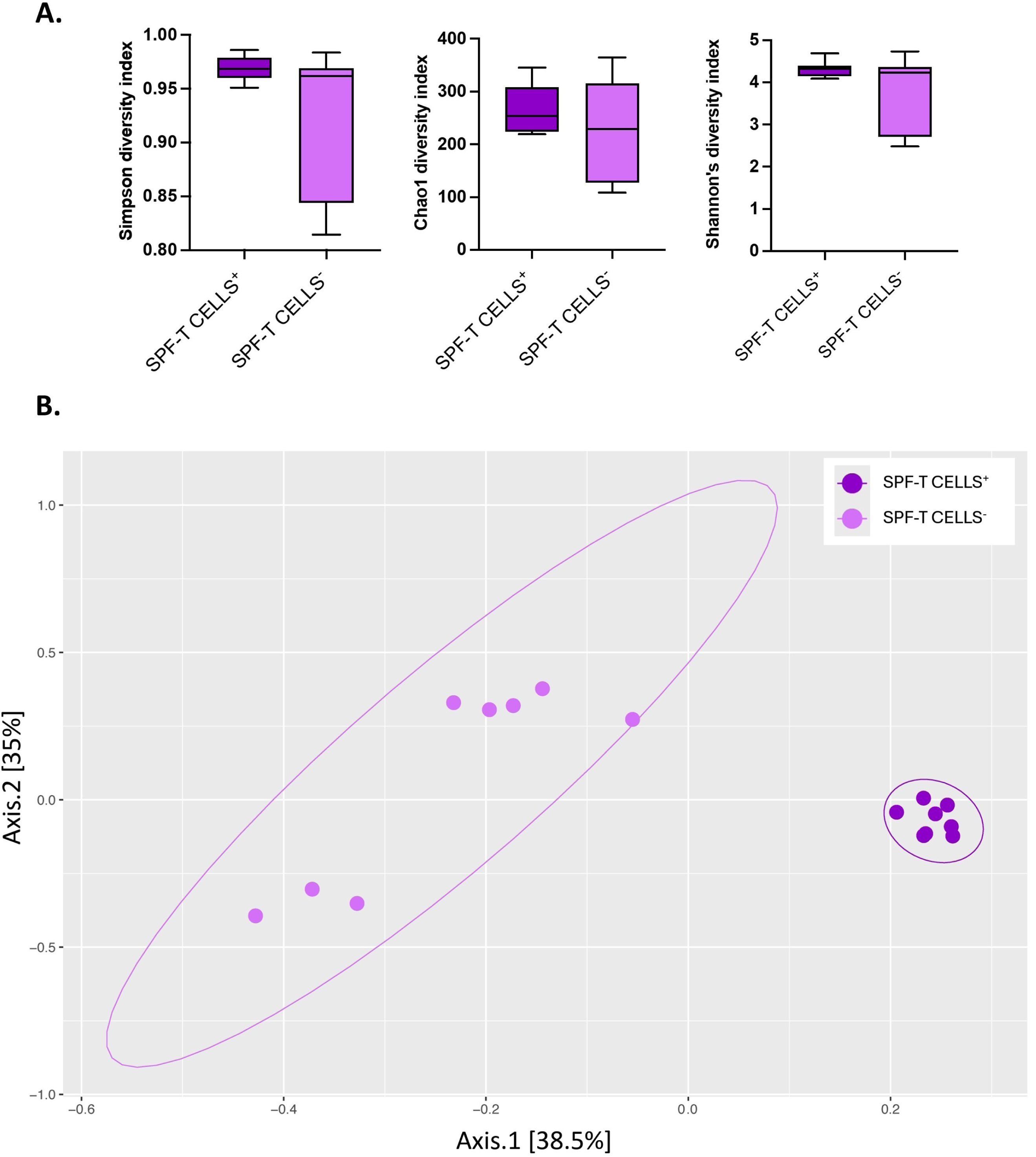
Microbiota profiles are similar in wildtype and T cell-deficient germ-free mice after SPF colonization. **A**. Alpha Diversity was measured in samples from wildtype germ-free mice colonized with SPF microbiota (SPF-T CELLS^+^) and T cell-deficient germ-free mice colonized with SPF microbiota (SPF-T CELLS^-^) groups using the Simpson, Chao1, and Shannon Diversity Indices. Data are expressed as mean ± SEM, n= 8 mice per group, analyzed using either a Student’s t-test (Chao1) or a Mann-Whitney test (Simpson, Shannon) based on the distribution of the data. There were no statistical differences between the groups. **B.** Beta diversity was measured by generating a Bray-Curtis dissimilarity matrix, data was then plotted using a principal coordinate analysis plot. PERMANOVA analysis was used for the Bray-Curtis Dissimilarity to compare SPF-T CELLS^+^ and SPF-T CELLS^-^ groups, which clustered separately (999 permutations, F=8.3226, p<0.001; N=8 mice/group). Each dot represents an individual animal.

We also measured SFB levels in the terminal ileum by qPCR in both normal and T cell-deficient germ-free mice colonized with SFB and SPF microbiota. Germ-free mice had no detectable SFB, while the SFB monocolonized mice showed significant enrichment of SFB compared to SFB levels in wildtype and T cell-deficient SPF colonized mice (Supplementary Figure 4). This result was expected, since SFB is only one member of the SPF microbiota, and the degree of colonization is likely not as great due to the presence of other competing bacteria. Interestingly, the quantity of SFB microbiota was significantly greater in T cell-deficient mice than wildtype mice colonized with SFB (Supplementary Figure 4). Together, these data reveal that the changes observed in the small intestinal transit and neuronal density were not likely the result of markedly different microbiota composition or the degree of colonization.

### Colonization of the ileum and colon induces Th17 differentiation in the intestinal lamina propria

To determine which T cell subsets were present in the intestinal lamina propria that might contribute to the changes in motility, enteric neuronal density, and enteric neurogenesis, we next assessed the impact of SFB and SPF colonization on the proportion of T cell subsets in the ileum and colon (Figure 6, Supplementary Figure 5). Th17 cells (CD3^+^CD4^+^RORγT^+^ cells) represented only a small fraction of CD4^+^ T cells in wildtype germ-free mice (Figure 6A-B). In the ileum, this increased ∼10 fold in mice colonized with SFB and ∼20 fold in mice with an SPF microbiota (Figure 6B). In the colon of germ-free mice, Th17 cells represented about 6% of the total CD4^+^ T cells and this doubled in SPF colonized mice. SFB colonization increased Th17 cells to about 10% of the total CD4^+^ cells. In contrast to the differentiation of Th17 cells by colonization, the proportions of regulatory T cells (CD3^+^CD4^+^FOXP3^+^) were significantly reduced after SFB and SPF colonization in the ileum, and after SPF colonization in the colon (Figure 6C)^36^. In contrast, the proportions of induced regulatory T cells expressing RORγT^+^ (CD3^+^CD4^+^FOXP3^+^RORγT^+^)^36^. were increased in the colon of mice colonized with SFB and SPF (Figure 6D). However, in the ileum, we only observed an increase in this population in mice colonized with SPF microbiota (Figure 6D). We next assessed the proportions of total innate lymphoid cells (ILCs) after colonization in the ileum or colon and observed no significant alterations in either region of the gut (Figure 6E). We then determined the frequencies of type 3 ILCs (ILC3), as they are a predominant population in the ileum and colon compared to other ILCs subsets and play a crucial role in contributing to gut homeostasis^39, 40^. We saw a small, but significant, reduction of ILC3s in the ileum of SPF colonized mice (Figure 6F). We also saw a significant increase in the frequency of ILC3 in the colon of SFB colonized mice, which was not observed in SPF colonized mice (Figure 6F).

**Figure 6.**
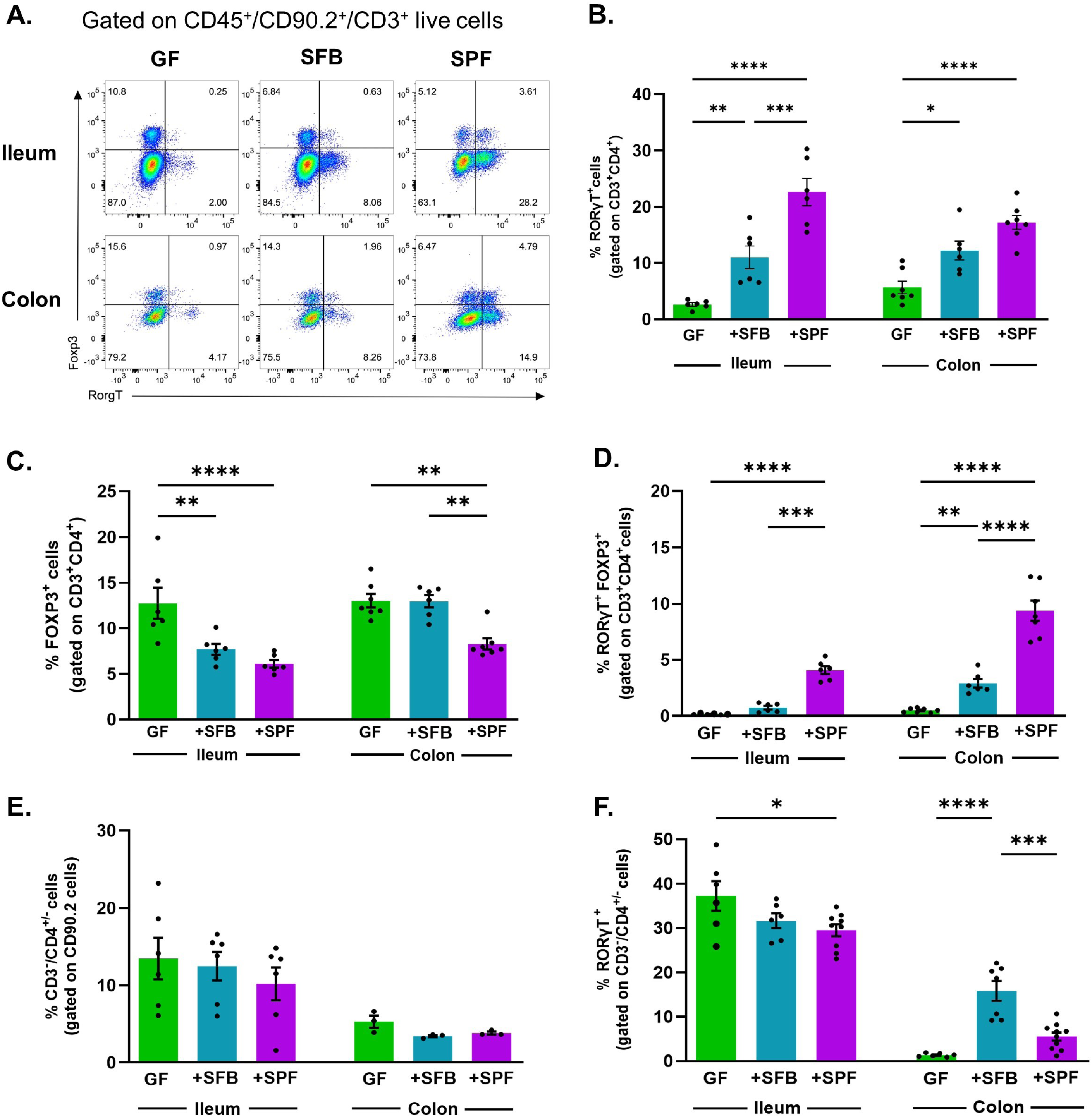
Microbiota colonization induces an expansion of Th17 cells and regulatory T cells expressing RORγT in the lamina propria of the ileum and colon. Germ-free mice (GF) were colonized with SFB or with SPF microbiota and small intestinal and large intestinal lamina propria cells were isolated. **A.** Representative dot plots showing the proportions of CD45^+^CD90.2^+^CD3^+^CD4^+^ cells expressing Foxp3 or RORγT in the small intestinal lamina propria (top panels) and colonic lamina propria (bottom panels) in GF mice, GF mice colonized with SFB and SPF microbiota. **B-D.** Proportions of RORγT Th17 cells (**B**), FoxP3^+^ regulatory T cells (**C**) and regulatory T cells expressing FoxP3 and RORγT (**D**) in the ileum and colon of GF, SFB and SPF colonized mice. All the parental populations were gated on CD45^+^CD90.2^+^ live cells. **E-F.** Proportions of ILCs (**E**) and ILC3 cells (**F**) gated on CD45^+^CD90.2^+^CD3^-^CD4^+/-^. Each dot represents an individual animal. Data are pooled from 2-4 independent experiments. Data were analyzed using two-way ANOVA, followed by Šídák’s multiple comparison test. N=3-8/group. * p<0.05, ** p<0.01, *** p<0.001, **** p<0.0001.

Staining lamina propria cells for the T cell receptor beta chain (TCRβ) confirmed the absence of T cells in the TCRβ^-^deficient mice (Supplementary Figure 6A). A similar pattern was noted for ILCs in T cell-deficient mice, with no changes observed in this cell population after colonization with SFB or SPF in either region of the gut in the absence of T cells, confirming that ILCs are developmentally independent of T cells (Supplementary Figure 6B)^41, 42^.

Together these data confirm that colonization of germ-free mice with SFB and SPF modulates the pattern of immune cells in the intestinal lamina propria^19, 36, 43^. These changes correlated with the increases in neuronal density and enteric neurogenesis observed after colonization with SFB and SPF microbiota.

### Colonization of the ileum and colon elevates intestinal cytokines

Given our findings on the impact of T cells on the ENS and the increased numbers of Th17 cells in the colonized mice, we examined the levels of IL-17A, and a panel of other cytokines and chemokines (Supplementary Tables 1 & 2). As expected, we found that in germ-free mice IL-17A was virtually undetectable in both the ileum and colon (Figure 7A, 7B). In the ileum, SFB monocolonization increased IL-17A levels, which were further elevated in animals colonized with SPF microbiota (Figure 7A). In the colon, SPF but not SFB colonization induced significantly higher levels of IL-17A compared to germ-free mice (Figure 7B). The levels of IL-17A in the ileum and colon of T cell-deficient mice were not elevated by colonization with either SFB or SPF microbiota (Figure 7A, B)

**Figure 7.**
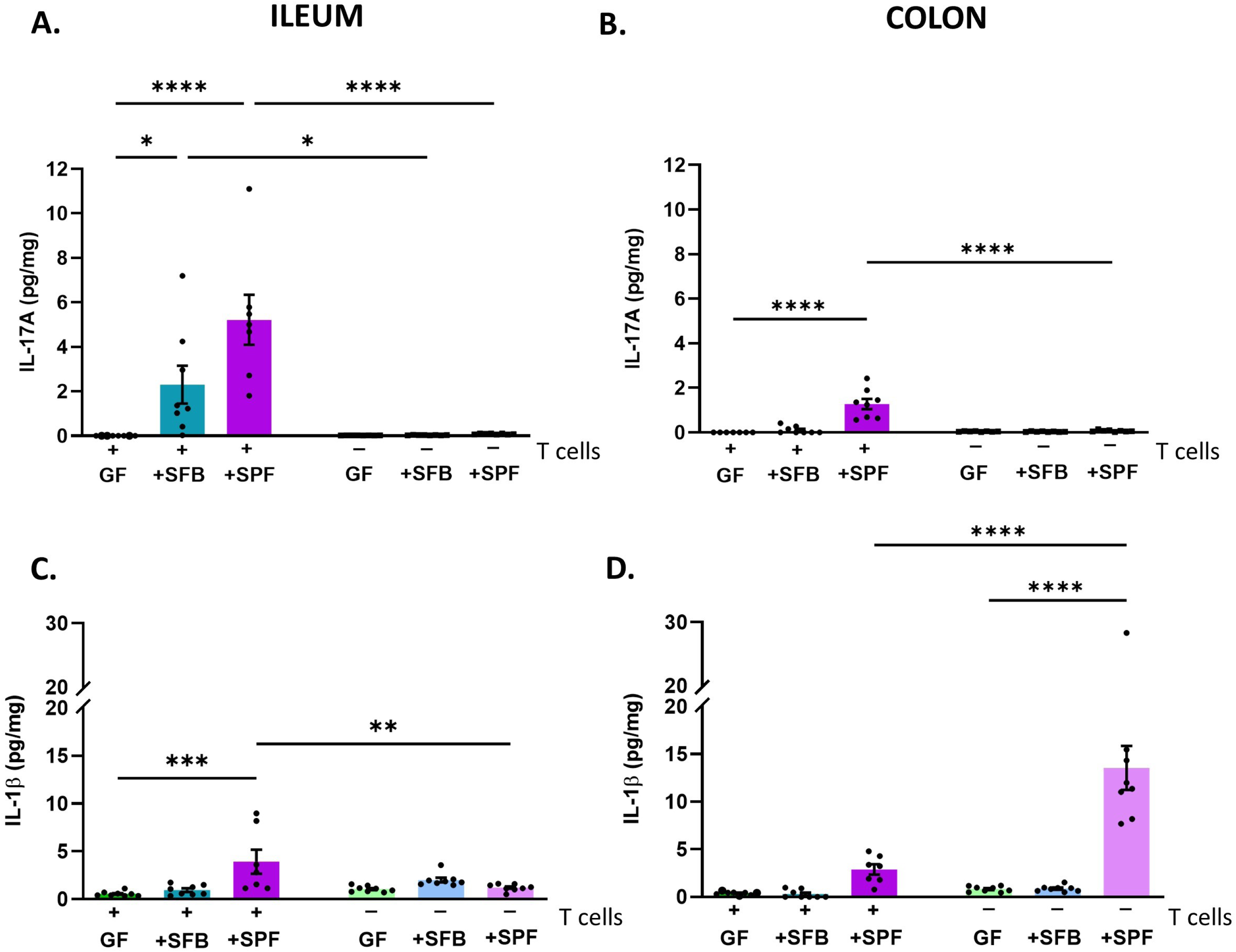
Microbiota colonization increases IL-17A and IL-1β in the ileum and colon in a T cell-dependent manner. **A-B**. IL-17A levels in the ileum (**A**) and colon (**B**) of wildtype germ-free mice (GF), GF mice colonized with SFB or SPF microbiota (+), T cell-deficient GF mice, and T cell-deficient GF mice colonized with SFB or SPF microbiota (-). **C-D.** IL-1β levels in the ileum (**C**) and colon (**D**) of wildtype germ-free mice (GF), GF mice colonized with SFB or SPF microbiota (+), T cell-deficient GF mice, and T cell-deficient GF mice colonized with SFB or SPF microbiota (-). Each dot represents an individual animal. Data are pooled from 2-4 independent experiments run for each genotype. Data were analyzed using two-way ANOVA, followed by Šídák’s multiple comparison test. N=7-8/group. * p<0.05, ** p<0.01, *** p<0.001, **** p<0.0001.

We assessed 16 other cytokines and chemokines and found that IL-1β showed a similar pattern of expression to IL-17A (Figure 7C-D, Supplementary Tables 1 & 2). Of note, the level of IL-1β in the colon was markedly stimulated by SPF colonization in T cell-deficient mice. It is interesting to note that the restoration of normal small intestinal transit correlated with the increased expression of IL-1β and IL-17A and the induction of Th17 cells in the small intestine.

### IL-17 receptor expression in the adult ENS

Single-cell RNA sequencing data suggested that the IL-17 receptor (*Il17a-e*) is expressed in the mouse ENS^25, 26^. We used immunohistochemistry to examine the expression of IL-17R in germ-free and colonized wildtype and T cell-deficient mice (Supplementary Figure 7). IL-17R appears to be expressed in most neurons in both plexuses and the levels of expression mirrored the changes in neuronal density.

### Immunoneutralization of IL-1β and IL-17A reduced neuronal density, Sox2 and nestin expression in SPF colonized mice, but had no effect on small intestinal transit

Given the correlation between IL-17A and IL-1β levels and Th17 cells with the normalization of small intestinal transit and induction of Sox2 in the myenteric plexus, we conducted studies using neutralizing antibodies for IL-1β and IL-17A in wildtype mice to determine if these cytokines were responsible for mediating these effects.

Wildtype germ-free mice were colonized with an SPF microbiota and administered IL-1β and IL-17A neutralizing antibodies 3 times/week for 4 weeks. Cytokine measurements confirmed that the neutralizing antibody treatment significantly reduced the levels of IL-1β and IL-17A in the ileum and IL-17A in the colon (Supplementary Figure 8). The levels of IL-1β levels were unchanged in the colon. Treatment of SPF mice with IL-1β and IL-17A neutralizing antibodies had no effect on any of the other cytokines and chemokines measured (Supplementary Table 3).

We then examined the effect of IL-1β and IL-17A neutralization on the gut microbiota composition and T cell populations in the intestinal lamina propria. We found no differences in either alpha or beta diversity of the neutralizing antibody-treated SPF-colonized germ-free mice when compared to isotype control SPF-colonized germ-free mice (Supplementary Figure 9A-B). Furthermore, the relative abundance of these groups shows a clear similarity in microbial composition among samples (Supplementary Figure 10). Similarly, immunoneutralization of IL-1β and IL-17A did not alter T cell populations (Supplementary Figure 11A-E) but did lead to a significant increase in the proportion of ILC3 cells in the colon (Supplementary Figure 11E).

No differences in the length of the ileum (38.9 ± 0.5cm Isotype control, 38.6 ± 0.5cm anti-IL-1β and IL-17A-treated [n=8]) was found between the isotype control and IL-1β and IL-17A antibody treated mice and treatment did not reverse the effect of SPF microbiota on small intestinal transit (Figure 8A).

**Figure 8.**
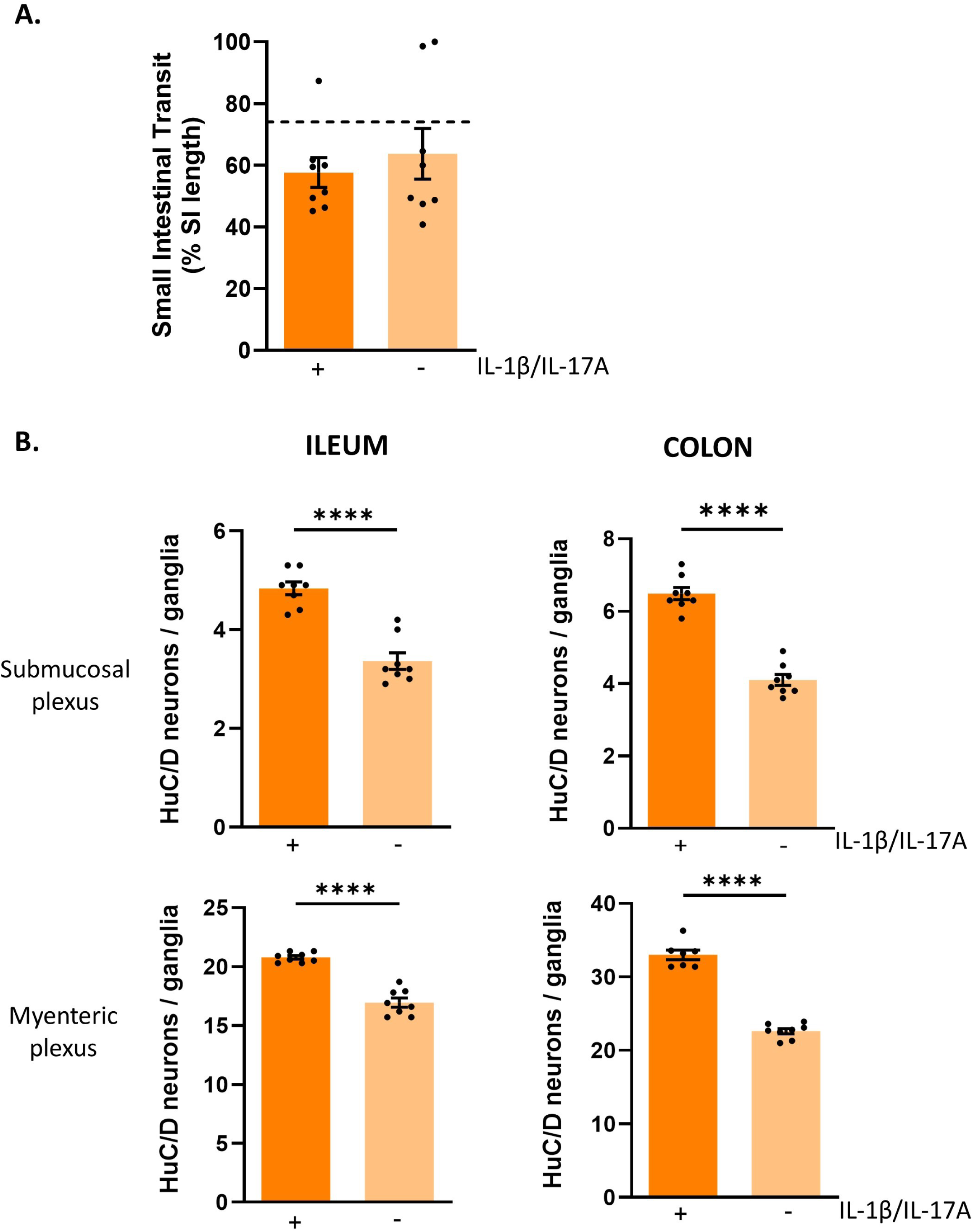
Immunoneutralization of IL-1β and IL-17A reduces neuronal density but has no effect on small intestinal transit. **A**. Small intestinal transit was measured after oral gavage of carmine red in wildtype SPF colonized germ-free mice treated with isotype control antibodies (+) or a mixture of IL-1β/IL-17A neutralizing antibodies (-). Each dot represents an individual animal. Dashed line indicates small intestinal transit measured in wildtype mice colonized with SPF from birth. Data were analyzed with a Student’s t test. N=8/group. **B.** Enteric neuronal density was assessed using counts of HuC/D neurons in the submucosal and myenteric plexuses in wildtype SPF colonized germ-free mice treated with isotype control antibodies (+) or a mixture of IL-1β/IL-17A neutralizing antibodies. Each dot represents an individual animal. Data are pooled from 2 independent experiments. Data were analyzed with a Student’s t test. N=8/group. **** p<0.0001.

In contrast, treatment of SPF mice with IL-1β and IL-17A neutralizing antibodies significantly reduced the SPF-increased neuronal density in the ENS to levels seen in germ-free mice (Figure 8B). Similarly, the expression of Sox2 in the myenteric plexus of the ileum and colon were significantly reduced by treatment with the neutralizing antibodies (Supplementary Figure 12). Like Sox2, nestin expression was also reduced in the myenteric plexus of the ileum and colon (Supplementary Figure 12). Together, these data suggest that intestinal cytokines IL-1β and IL-17A and T cells are important mediators of the neurogenic response of the ENS to the presence of the gut microbiota. Whilst T cells are important players in the microbiota ENS axis, mediators other than IL-1β and IL-17A regulate small intestinal motility after colonization of the gut of germ-free mice.

## Discussion

In this study we showed that in adult mice, T cells are important for the normalization of intestinal motility in germ-free mice colonized with an SPF microbiota, but the cytokines IL-1β and IL-17A are not the molecular mediators, since immunoneutralization was unable to block the response to colonization. Monocolonization of germ-free mice with SFB was not sufficient to normalize intestinal motility. Four weeks of colonization of the gut by SFB and SPF microbiota markedly increased Th17 cells and Treg expressing RORγ^+^T cells in both the ileum and colon of wildtype GF mice and restored neuronal density to control levels. Restoration of neuronal density was largely T cell-independent, but in contrast to motility, was mediated by the cytokines IL-1β and IL-17A. The exception to this was the restoration of neuronal density in the ileum in response to SFB, which was T cell-dependent. In addition to showing that the composition of the microbiota altered the mucosal immune environment, we also showed that T cell-deficient mice had an altered cecal microbiota composition, furthering the concept of bidirectional regulation of the gut microbiota by the immune system^44, 45^.

Sox2 expression in myenteric neurons has been shown to be associated with neurogenesis^28, 30^. Here we showed that in the myenteric plexus, colonization with SFB and SPF microbiota enhances the expression of Sox2 in a T cell-dependent manner in the ileum, but independently of T cells in the colon. The enhanced expression of Sox2 in response to SPF microbiota is mediated by the cytokines IL-1β and IL-17A. Both SFB and SPF stimulated nestin expression in the myenteric plexus of the ileum of wildtype mice, where the levels were higher in T cell-deficient mice, but there was no evidence that colonization altered the levels of expression. In the colon, nestin expression was stimulated by SFB in wildtype mice, and as in the ileum, there was no evidence that colonization altered the levels of expression in T cell-deficient mice. This appears to be the first study to show nestin expression in the ENS is regulated by T cells.

Adult mice born and raised germ-free, or SPF mice treated with antibiotics to deplete gut bacteria, have slower GI transit than conventional SPF mice^8, 9, 11, 12, 15, 16, 17, 18, 21, 46^. Motility is restored when adult germ-free mice are colonized with a mouse SPF microbiota^15^, human fecal microbiota^16^, indigenous spore-forming bacteria^18^, *E. coli*^11^, *L. rhamnosus*^11^ or when antibiotic treatment is stopped^8, 11^. There are persistent changes to motility when antibiotics are given to neonates and the mice are studied as adults^46^. The continuous presence of microbiota is essential for normal gut motility, since mice transiently colonized revert to a slowed transit once the bacteria are cleared^11^. We confirmed that SPF microbiota enhanced small intestinal transit to levels similar to that of conventional SPF mice, but found that monocolonization with SFB was insufficient, despite its ability to restore neuronal density.

Interestingly, the effects of SPF colonization were lost in T cell-deficient mice, implicating T cells in the regulation of intestinal motility by the microbiota for the first time to our knowledge. The gut microbiota regulates motility by controlling the excitability of intrinsic primary afferent neurons (AH cells)^47^, enhancing serotonergic signaling pathways^15, 16, 18^, and/or toll-like receptor signaling^11, 34, 48^, though the exact mechanisms in different regions of the gut remain to be fully elucidated. How T cells are mediating their effects has not yet been determined. We hypothesized that IL-1β and IL-17A were involved in the microbiota-mediated T-cell regulation of motility based on the correlation of their elevated levels with the enhanced transit. However, immunoneutralization did not alter the normalized intestinal transit following colonization with an SPF microbiota, suggesting other mediators are involved in this mechanism.

Administration of antibiotics to adult mice reduces the density of neurons in the myenteric plexus of the ileum^8, 12, 14^ and colon^8, 14, 34, 46, 48, 49^, but not the duodenum^14, 49^. Cessation of antibiotics leads to a recovery of neuronal density in the small and large intestines^8, 14, 34^. The submucosal plexus has been less intensively investigated^8, 46^, but the findings are similar. These data suggest that the microbiota is essential for the maintenance of enteric neuronal homeostasis and that the ENS has an intrinsic neurogenic capacity in adulthood. Studies in adult germ-free mice are generally consistent with these findings, suggesting that the capacity to respond to the gut microbiota is intrinsic to the ENS, even if it is not exposed to a microbiota during the neonatal period. Thus, there is a reduced density of myenteric neurons in the jejunum^20^, ileum^14, 20^ and colon^34, 50^ reported by most studies. However, there are exceptions in the literature where either there was no change in myenteric neuronal density in the colon^14^ or that there was an increased neuronal density in the colon of germ-free mice^19^. There is no obvious explanation for the differences between these studies. When germ-free mice were colonized with altered Schaedler flora^20^, SPF microbiota^14^, or *B. thetaiotaomicron*^50^, or treated with the TLR2 agonist lipoteichoic acid^34^ neuronal density was increased. In the study where neuronal density was increased in germ-free mice, colonization with *C. ramosum* or *P. magnus*, reduced it^19^. In our study, we asked whether the potent inducer of Th17 cells, SFB, would alter adult enteric neurogenesis, since it did not restore motility. We found that SFB was able to fully restore neuronal density in the ENS of wildtype mice. Interestingly, in the ileum, but not the colon, this was dependent on T cells. These results suggest there are different mechanisms involved in the response to SFB in the ileum (where it colonizes), vs the colon. In the myenteric plexus, Sox2-expressing enteric neurons were stimulated by SFB and SPF microbiota and this effect was lost in the ileum of T cell-deficient mice. These data suggest that the neurogenic response to SFB requires T cells in the ileum, but that other cell types (e.g., ILCs) mediate the response to SPF microbiota in that region of the gut. In the myenteric plexus of the colon the neurogenic response is T cell-independent.

Nestin has been proposed as a marker of enteric neurogenesis in the adult ENS^27, 31, 34, 35^. Nestin expression is found in healthy adult myenteric neurons, suggesting that there is a balance between cell proliferation and cell death under physiological conditions. The levels of nestin expression in the myenteric plexus of the colon observed in this study are similar to that previously reported^34^. Nestin expression was modulated by the gut microbiota, and interestingly, T cells are important in regulating the levels of nestin expression, since in T cell-deficient mice, levels increased. Whether this was due to the altered composition of the gut microbiota in these mice or the absence of T cells needs further investigation. We found no evidence of nestin expression in the submucosal plexus, and little Sox2 expression, so further work is needed to better define the markers of adult neurogenesis in this plexus. Nevertheless, throughout the gut, the neurogenic responses ultimately involves IL-1β and IL-17A, since immunoneutralization of these cytokines reduced the increase in neuronal density, Sox2 and nestin expression in enteric neurons in both enteric plexuses. These findings thus provide new insights into the mechanisms of the microbiota-neuroimmune dialogue that regulates intestinal physiology.

One defining characteristic of adaptive immune responses to the microbiota is that their development and establishment occurs in the complete absence of inflammation, a process that is referred to as homeostatic immunity^51, 52^. We have observed that T cells play a pivotal role in regulating gut motility. These adaptive responses are fundamental in containing the microbiota in its physiological niche and highly entwined with tissue physiology contributing to the maintenance of tissue integrity and regulation as a whole^51, 52^. Along similar lines, Geuking *et al*. showed the colonization with a benign physiological defined microbiota, altered Schaedler flora, resulted in the compartmentalized activation and induction of intestinal Treg cells essential for successful establishment of intestinal CD4^+^ T cell homeostasis^53^. In our work we lost the normalization of intestinal motility in T cell-deficient germ-free mice colonized with SPF microbiota, and this effect was independent of IL-1β and IL-17A. These data are interesting in the light of observations that IL-17A in the context of sepsis and intestinal inflammation is associated with intestinal dysmotility, albeit indirectly, via the activation of macrophages^54, 55^. Whilst, T cells have long been implicated in the regulation of intestinal contractility in nematode infection^56^, our data shows that they are also importantly involved in the response to microbiota colonization. Further studies are needed to identify the mediators of this effect.

In conclusion, we show that T cells are important for the normalization of intestinal motility after colonization and that the cytokines IL-1β and IL-17A mediate the enteric neurogenic response to an SPF microbiota (Supplementary Figures 13 and 14). SFB monocolonization stimulates enteric neurogenesis, requiring T cells in the small intestine, but not the colon, but SFB alone is not sufficient to normalize small intestinal motility. Together, our findings provide new insights into the microbiota-neuroimmune dialogue that regulates intestinal physiology^57, 58^, highlighting a role for T cells in regulation of intestinal motility and a role for the cytokines IL-1β and IL-17A in enteric neurogenesis that occurs in adult mice in response to microbial colonization of the gut. And whilst this study focused on the impact of the intestinal microbiota, and T cells on the ENS and intestinal motility, it is important to recognize that these interactions are bidirectional, as the composition of gut microbiota is shaped by the ENS^7^, which in turns regulates intestinal physiology.

## Methods

### Animals

Germ-free male C57Bl/6 mice (10-13 weeks of age) and TCRβ^-^Tcrδ^-^ (B6.129P2-*Tcrb^tm1Mom^ Tcrd^tm1Mom^*/J)^59^ male mice (13-23 weeks of age) were provided by the University of Calgary International Microbiome Centre (IMC). Mice were either left untreated (germ-free) or were colonized with segmented filamentous bacteria (SFB) from birth (born to SFB breeding pairs) or were colonized with specific pathogen-free (SPF) microbiota for 4.1 ± 0.3 weeks by gavage with SPF microbiota. In brief, the cecal, small intestinal and colonic contents were collected from 2-3 SPF mice in an anaerobic chamber, homogenized in degassed phosphate buffered saline (PBS; 1:15 dilution), and gavaged into germ-free mice. Germ-free male TCRβ^-^ mice were either untreated (germ free), SFB-colonized by co-housing with an SFB mouse for 5.1 ± 0.2 weeks or colonized with SPF microbiota for 3.4 ± 0.1 weeks by gavage of SPF microbiota as above. All gnotobiotic mice were bred and maintained in flexible film isolators until experimental use, maintained at 22 ± 2°C on a 12-hour light-dark cycle with free access to sterilized food and water in the IMC where germ-free status is routinely monitored by culture-dependent and independent methods. Conventional male C57BL/6 SPF mice (11-12 weeks of age), from the same colony used to collect the gavage material for the SPF gavages described above, were housed in the animal facility at the University of Calgary at 22 ± 2°C on a 12-hour light-dark cycle. Mice had free access to sterilized food and water. Experimental mice were maintained in IsoPositive Cages (Tecniplast) until experimental endpoint. All procedures involving animals were approved by the University of Calgary Health Sciences Animal Care Commitee (AC19-0124, AC23-0135, AC21-0015, AC21-0051) and are in accordance with the guidelines established by the Canadian Council on Animal Care. Male mice were used based on our past observations where there were no differences in response to recolonization in male and female mice^8^.

### Neutralizing antibody treatment

All antibodies were obtained from BioXCell (Lebanon, NH, USA). Anti-mouse/rat IL-1β (BE0246) and anti-mouse IL-17A (BP0173) neutralizing antibodies (100mg/100ml each) or their isotype controls (polyclonal Armenian hamster IgG [BE0091] and mouse IgG1 [BP0083], respectively) were administered intraperitoneally 3 days following SPF microbiota gavage to germ-free SPF-colonized mice, 3 times/week for 4 weeks under sterile conditions. Antibodies were determined by the manufacturers to be endotoxin free (<2EU/mg). Dilutions were made up fresh from stock solutions kept at 4°C on the day of injection with dilution buffers provided by BioXCell (IP0065 and IP0070).

### Tissue collection

Mice were brought to the lab, weighed, and allowed to acclimate for 1 hour prior to assessment of small intestinal transit as described below. Following isoflurane anesthesia and euthanasia by cervical dislocation the entire gut was removed, ileum and colon length determined, and small intestinal transit quantified. Tissues were allocated for different analyses. A sample of terminal ileum (0.5cm) was collected for SFB quantification and samples of both the terminal ileum and proximal colon were collected for tissue cytokines (1 cm each) and immunohistochemistry (2 cm each). Caecal mater was collected for determination of bacterial composition. The remaining ileum and colon were used for flow cytometric analysis (see below).

### Small intestinal transit

Mice were gavaged with 250 μl of the non-absorbable dye carmine red (6% suspension in 0.5% carboxymethylcellulose in dH_2_O), and 30 min later, euthanized under isoflurane anesthesia. The small intestine was immediately removed and measured (pyloric sphincter to ileal–cecal junction) and the distance travelled by the dye front was also measured. Results are expressed as a percentage of the length covered by the dye over the total small intestinal length.

### Tissue cytokines

T-cell specific cytokines were measured in the terminal ileum and proximal colon collected following euthanasia. Tissues were homogenized following the instructions for the Pierce BCA-Protein Assay (ThermoFisher Scientific; Mississauga, ON, Canada; Cat #23225) and protein concentration in mg/ml tissue homogenate was determined. A multiplex panel of cytokines (Mouse High Sensitivity 18 Plex Discovery Assay; MDHSTC18) was run by Eve Technologies (Calgary, Alberta, Canada) and cytokine levels were expressed as pg/mg protein.

### Isolation of ileal and colonic lamina propria cells

Lamina propria cells were extracted from the ileum and colon using approaches previously published^21^. Briefly, the tissues were removed after euthanasia. The attached mesenteric fat and Peyer’s Patches were removed. After opening the tissue longitudinally, small pieces of approximately 2 cm were cut and transferred into epithelial cell removal solution (1x HBSS, 5% inactivated fetal calf serum (iFCS), 10 mM HEPES and 5 mM EDTA) and incubated on a shaker at 37°C for 20 min, repeated twice. The tissue was minced prior to digestion in supplemented RPMI (Collagenase type VIII (Sigma-Aldrich, St. Louis, MO, USA), DNAse I 10U/mL (Roche, Indianapolis, IN, USA), and 5% iFCS at 37°C for 20 min. Digestion was deactivated using cold RPMI containing 10% iFCS. The tissue was further dissociated by manual suction using a serological pipette and filtered through a 100 μm cell strainer, spun down and washed twice with FACS buffer (1x PBS, 2% iFCS, 2 mM EDTA). The cell suspensions were used for flow cytometry.

### Flow cytometry

Isolated ileal and colonic lamina propria cells were blocked using anti-CD16/32 antibody (clone 2.4G2; BD Biosciences, Mississauga, ON, Canada) for 20 min. Then, cells were stained for 30 min with antibodies for specific markers. Nonviable cells were identified using viability BV510 (clone FVS510 BD Biosciences) and sequentially stained with the following surface antibody mixes: CD45 PE-Cy7 (clone 30F11BD BD Biosciences), CD90.2 Alexa 700 (clone 53-2.1 Biolegend, San Diego, CA, USA), CD3 APC-Cy7 (clone 17A2 BD Biosciences) and CD4 FITC (clone GK-1.5; BD Biosciences), TCRβ-PERCP-Cy5.5 (BD Biosciences). Intracellular staining was performed using the Transcription Factor Staining Buffer Set (BD Biosciences), as well as the monoclonal antibodies RORγT BV421 (clone Q3-378; BD Biosciences), Foxp3 AF647 (clone MF23 BD Biosciences), all at a dilution of 1:100, following the instructions of the manufacturer. Samples were acquired using the flow cytometer FACS Canto II (BD Biosciences) and the corresponding fluorescence minus one (FMO) staining was used as a control. Data was analyzed using FlowJo software (Tree Star Inc., Ashland, OR, USA). The gating strategies used to determine the cell types are shown in Supplemental Figure 5. Briefly, for the gating strategy, we first gated singlets for aggregate exclusion. After that, for all analyses, we used CD45^+^BV510^-^ to discriminate viable cells. FSC vs SSC control was used to discriminate lymphocytes. CD90^+^ was used for lymphoid cells followed by CD4^+^CD3^+^ as a marker for T cells. We then used transcription factors to differentiate our target cell populations: for Th17, we used RORγ^+^; iTregs, RORγ^+^FOXP3^+^; Tregs, FOXP3^+^; Total Tregs. For ILC3, we used RORγT^+^ cells in CD4^-^CD3^-^ cells. Data are presented as the frequency’s gated on the parent population, as indicated in the Figures and Figure legends.

### Immunohistochemistry and neuronal quantification

Dissection of submucosal and myenteric plexuses were done similarly for the ileum and colon as previously published^8^. Tissue samples for wholemount preparations were initially placed in PBS containing 1 μM nifedipine (Sigma-Aldrich) for 10 min. Tissues were then opened along the mesenteric border and pinned out in a Petri dish containing Sylgard coating with the mucosal side facing up. Samples were fixed in Zamboni’s fixative for 24 hours at 4°C for labeling with anti-HuC/D and anti-Sox2, or 4% paraformaldehyde (ThermoFisher Scientific, Waltham, MA, USA) for 2 hours at 4°C for labeling with anti-HuC/D and anti-IL-17R or anti-nestin. Fixed tissues were then washed with PBS containing 0.01% sodium azide (3 times, 10 min each) and stored at 4°C until further processing. Submucosal and myenteric preparations were obtained by gently scraping the mucosa away and peeling off the submucosal plexus, and the myenteric preparations were made by stripping off the mucosa/submucosa layers and the circular muscle leaving behind a preparation consisting of the longitudinal muscle and adherent myenteric plexus.

Samples were then processed for immunohistochemistry. When using anti-Sox2 antibodies, samples were pre-treated with DMSO (VWR International, Edmonton, AB, Canada) for 30 min at room temperature prior to primary antibody incubation. Tissues were incubated with primary antibody (Table 1) for 48 h at 4°C. Samples were washed (3 times, 5 min each) with PBS containing 0.1% Triton X100 (Sigma-Aldrich), and then incubated with secondary antibody (Table 1) for 1–2 h at room temperature followed by washing (3 times, 5 min each) with PBS+Triton X100. Double labeling was done sequentially, so samples were washed with PBS+Triton X100 (3 times, 10 min each), and the second labeling was performed following the same protocol. Whole-mount preparations were mounted with bicarbonate-buffered glycerol on adherent microscope slides and stored in the fridge under dark conditions until analyses were performed. Antibodies used were diluted in a PBS solution containing 0.1% Triton X-100, 0.1% bovine serum albumin, 0.05% sodium azide, 0.04% EDTA.

Table 1.

**Table 1.**
| **Antibody** | **Dilution** | **Source** | **Identifier** |
| --- | --- | --- | --- |
| Mouse anti-HuC/D | 1:200 | Invitrogen | Cat# A21271; RRID: AB_221448 |
| Rabbit anti-HuC/D | 1:200 | Abcam | Cat# ab210554; RRID: AB_210554 |
| Goat anti-Sox2 | 1:200 | R&D Systems | Cat# AF2018; RRID: AB_355110 |
| Mouse anti-Nestin | 1:100 | Abcam | Cat# ab6142; RRID: AB_305313 |
| Goat anti-IL17R | 1:100 | R&D Systems | Cat# AF448; RRID: AB_355366 |
| Donkey anti-mouse AF488 | 1:200 | Jackson ImmunoResearch | Cat# 715-546-151; RRID: AB_2340850 |
| Donkey anti-rabbit AF488 | 1:200 | Jackson ImmunoResearch | Cat# 711-545-152; RRID: AB_2313584 |
| Donkey anti-goat CY3 | 1:200 | Jackson ImmunoResearch | Cat# 705-165-147; RRID: AB_2307351 |
| Donkey anti-mouse CY3 | 1:200 | Jackson ImmunoResearch | Cat# 715-165-150; RRID: AB_2315777 |

Quantification of the specifically labeled markers (HuC/D, Sox2, Nestin, IL17R) was performed using a Zeiss Axioplan fluorescence microscope (Zeiss Canada, Toronto, ON, Canada). Ten-15 submucosal or myenteric ganglia were randomly selected, and all cells counted, these counts were then averaged and data for each animal is presented. All quantification was done by an experimenter blinded to the treatment group.

### SFB quantification

Ileal samples were weighed and whole genomic DNA was extracted using the PowerFecal Pro® DNA Isolation Kit as per the manufacturer’s instructions (Qiagen, Germantown, MD, USA). 1 μl of each sample was used for 2-step real-time PCR amplification using SFB-specific primers (SFB736F: GACGCTGAGGCATGAGAGCAT, SFB844R: GACGGCACGGATTGTTATTCA)^60^ and PerfeCTa SYBR Green SuperMix (QuantaBio, Beverly, MA, USA). The annealing temperature was 60°C and a melting curve was generated at the end of the cycles.

### Bacterial composition and analysis

Cecal samples were collected at the time of sacrifice and stored at –80°C until DNA isolation. Bacterial DNA was isolated from samples using the QIAamp PowerFecal Pro DNA Kit (Qiagen, 51804). The V4 hypervariable region of the bacterial 16S rRNA gene was amplified by using barcoded primers: (16SV4Fwd: AATGATACGGCGACCACCGA BARCODE TATG-GTAATTGTGTGCCAGCMGCCGCGGTAA and 16SV4Rev: CAAGCAGAAGACGGCATAC-GAGAT BARCODE AGTCAGTCAGCCGGACTACHVGGGTWTCTAAT) (Integrated DNA Technologies, Coralville, IA, USA). KAPA HiFi polymerase (Roche) was used for the reactions with the following cycling conditions: initial denaturation at 98°C for 2 min, 25 cycles of 98°C for 30s, annealing at 55°C for 30s, extension at 72°C for 20s and final elongation at 72°C for 7 min. Following amplification, individual PCR libraries were pooled, and the concentration of each sample was measured using a Qubit fluorometer (ThermoFisher, Waltham, MA, USA). The 16S rRNA V4 gene amplicon sequencing was performed with a V2-500 cycle cartridge on the MiSeq platiorm (Illumina, San Diego, CA, USA). The raw fastq files were processed with the DADA2 pipeline 1.16 (R package, version 1.30.0). The quality profiles of the forward and reverse reads were visualized, and the reads were trimmed to remove the V4 primer regions and lower-quality reads. The forward and reverse reads were then merged, and an amplicon sequence variant table (ASV) table was generated. A taxonomic classification was assigned to each ASV using the SILVA ribosomal RNA gene database project (version 138.1)^61^. Next, the phyloseq package was used to create a phyloseq object using the DADA2 outputs (R package, version 1.46.0). This phyloseq object was used for all further downstream analysis. Alpha diversity was measured using the Shannon, Chao1 and Simpson diversity indices. Beta diversity was analyzed using the Bray-Curtis dissimilarity matrix which was visualized using a Principal Coordinate Analysis (PCoA) plot. Relative Abundance was measured by examining the top thirty most prevalent ASVs in each sample. Finally, differential abundance between groups was measured using DESeq2 (R package, version 1.42.0). The alpha value was set to p<0.01 to decrease the false discovery rate (FDR) and significant changes in ASV abundances were ploted on a logarithmic scale base 2-fold change. The following statistical tests were carried out for the analyses, Student’s t test (alpha diversity), PERMANOVA (beta diversity), and Wald test (differential abundance).

### Data presentation and analysis

Data were graphed and analyzed using GraphPad Prism 10 (GraphPad Sotiware, La Jolla, Ca, USA) and are presented as the mean ± standard error of the mean (SEM). Microbiota data was graphed and analyzed using R (R package, version 1.42.0). Data were tested for normality using the Kolmogorov-Smirnov test. Subsequently, Student’s *t* test was used for analysis between two groups. For multiple comparisons a one-way ANOVA or two-way ANOVA, followed by Šídák’s multiple comparison test, as appropriate. The specific statistical test used is described in each figure legend as well as the group size. P < 0.05 was accepted as statistically significant.

## Acknowledgements

We are grateful to Karen Poon for assistance with the flow cytometry and Spencer Abbott for statistical analysis. We thank the University of Calgary Snyder Institute Bioinstrumentation lab for infrastructure support for qPCR, the Snyder Institute Nicole Perkins Microbial Communities Core Lab for flow cytometry support and the International Microbiome Facility for providing the germ-free animals.

## Author Contributions

PRMS, CMK, MDF, FAV, KDM, and KAS designed the studies; PRMS, CMK, LEW, MDF, and CO conducted the experiments and performed experimental data analyses; YBH performed the microbiota sequencing and bioinformatic data analyses; KDM and KAS provided study oversight and supervision; PRMS, CMK, LEW, FAV and KAS drafted the manuscript. All authors had full access to the study data and critically reviewed and approved the final manuscript for submission. KDM and KAS obtained funding for the study.

## ORCiD

PRMS. ORCID ID: 0000-0002-0216-7811

KDM. ORCID ID: 0000-0002-3900-9227

KAS. ORCiD ID: 0000-0001-9560-1711

## Supplementary Figure Legends

**Supplementary Figure 1.**
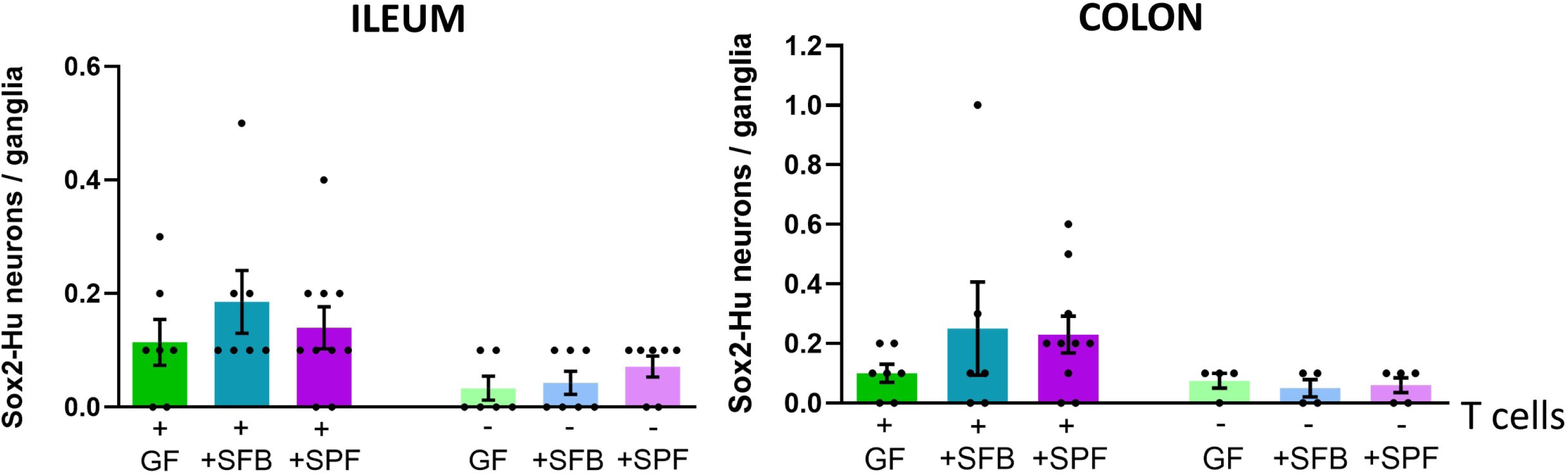
Sox2 is minimally expressed in submucosal neurons. Sox2-HuC/D neurons in the submucosal plexus of wildtype germ-free mice (GF), GF mice colonized with SFB or SPF microbiota (+), T cell-deficient GF mice, and T cell-deficient GF mice colonized with SFB or SPF microbiota (-) in the ileum (left panel) and colon (right panel). Each dot represents an individual animal. Data are pooled from 2-4 independent experiments run for each genotype. Data were analyzed using two-way ANOVA, followed by Šídák’s multiple comparison test. N=4-8/group. There were no statistical differences between the groups.

**Supplementary Figure 2.**
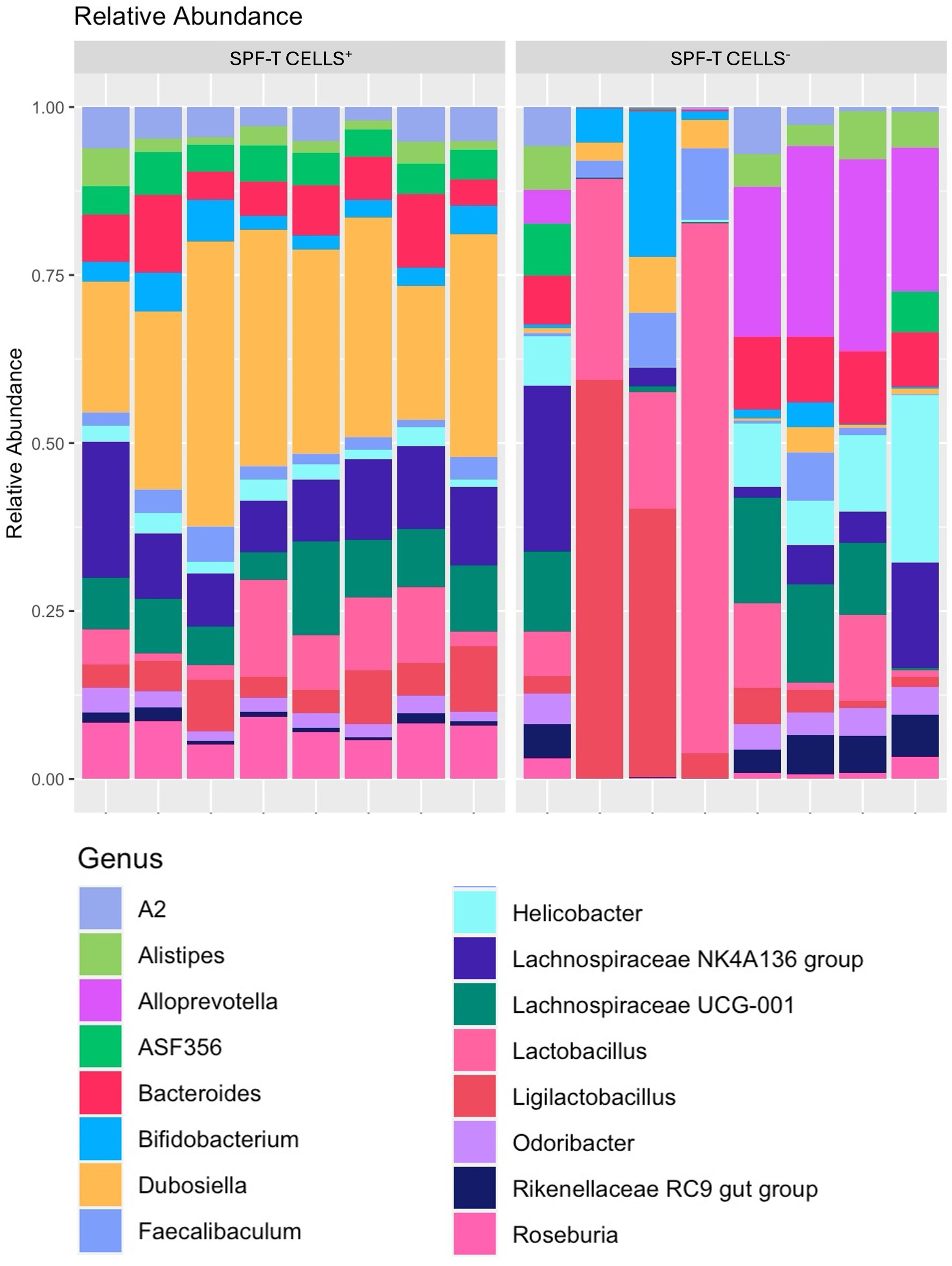
Top 30 prevalent amplicon sequencing variants (ASV) of SPF colonized mice. The top 30 prevalent ASVs were retrieved from wildtype germ-free mice colonized with SPF microbiota (SPF-T CELLS^+^) and T cell-deficient germ-free mice colonized with SPF microbiota (SPF-T CELLS^-^) and the relative abundance at the genus level was plotted per mouse.

**Supplementary Figure 3.**
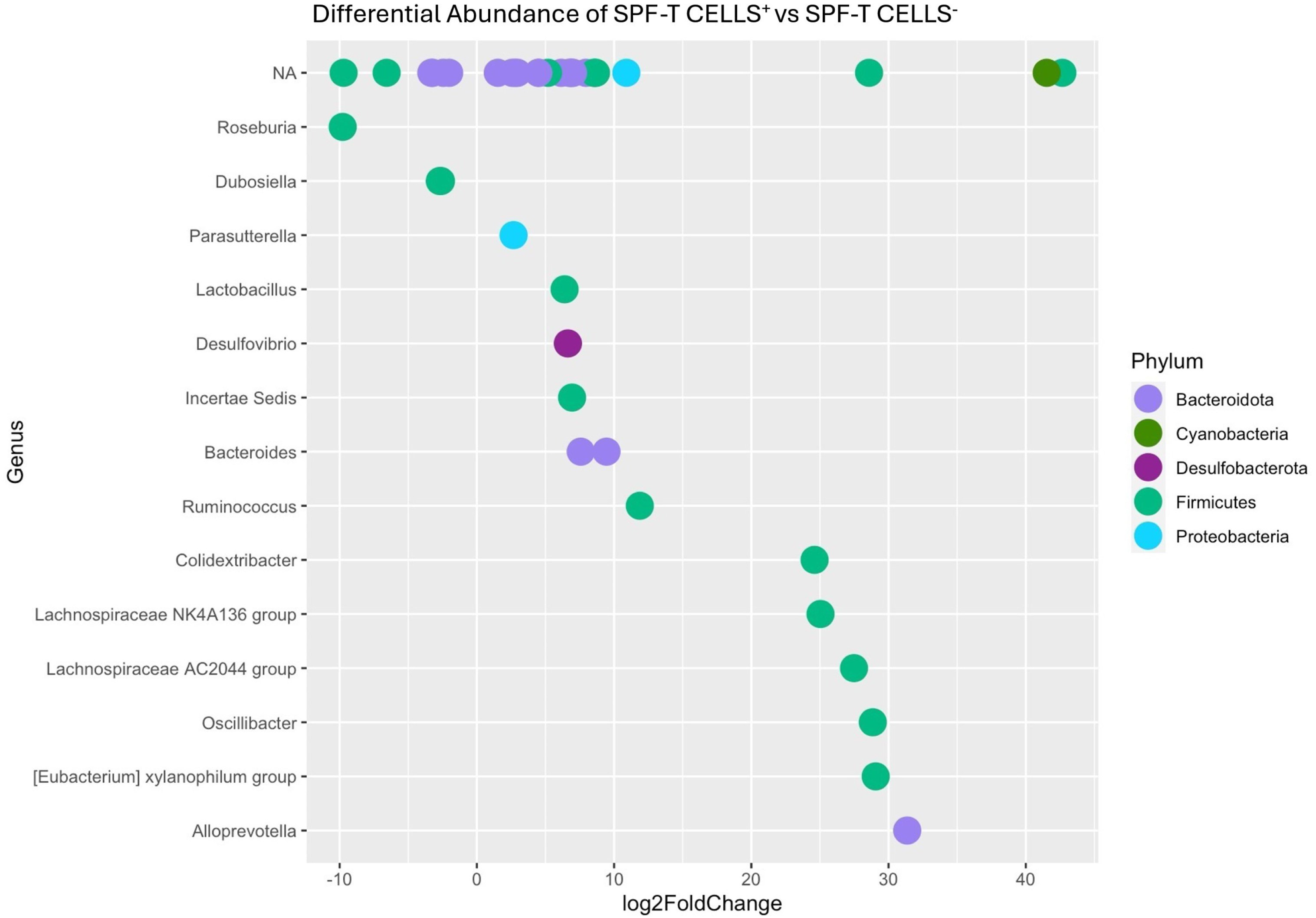
Differential abundance of amplicon sequencing variants (ASV) of SPF colonized mice. Log2 fold change of which ASVs were significantly increased (right) and decreased (left) in T cell-deficient germ-free mice colonized with SPF microbiota (SPF-T CELLS^-^) mice relative to wildtype germ-free mice colonized with SPF microbiota (SPF-T CELLS^+^) mice. Following colonization, T cell-deficient mice showed a significant decrease in *Roseburia* and *Dubosiella*, when compared to wildtype mice. Each point on the plot represents an ASV that is categorized at the genus level. ASVs were considered differentially abundant and plotted if the adjusted p-value was below p<0.01. DESeq2 analysis using the Wald Test was performed.

**Supplementary Figure 4.**
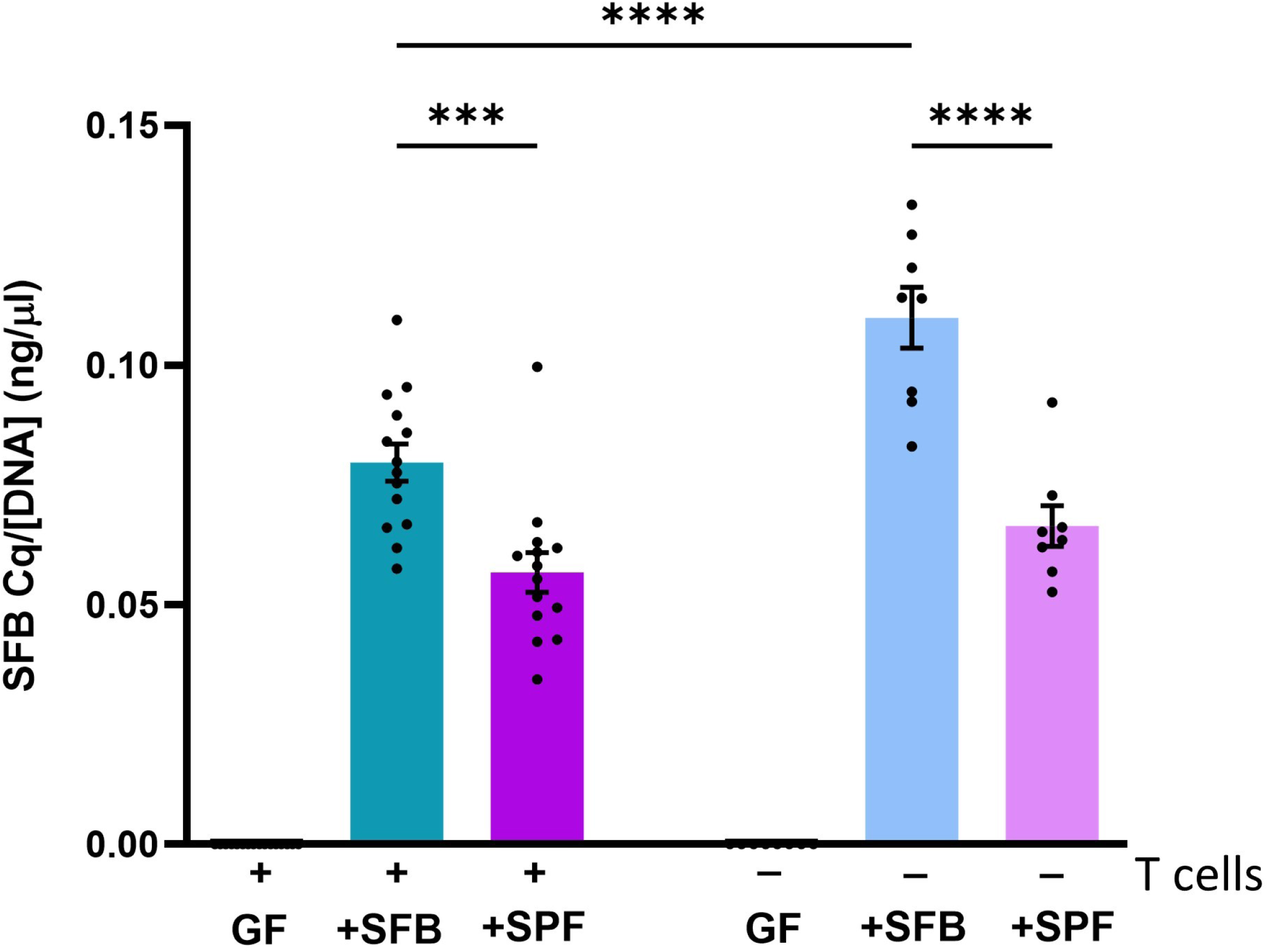
Quantification of SFB. SFB levels were quantified in ileal samples from wildtype germ-free mice (GF), GF mice colonized with SFB or SPF microbiota (+), T cell-deficient GF mice, and T cell-deficient GF mice colonized with SFB or SPF microbiota (-). Each dot represents an individual animal. Data are pooled from 2-4 independent experiments run for each genotype. Data were analyzed using two-way ANOVA, followed by Šídák’s multiple comparison test. N=8-18/group. * p<0.05, **** p<0.0001.

**Supplementary Figure 5.**
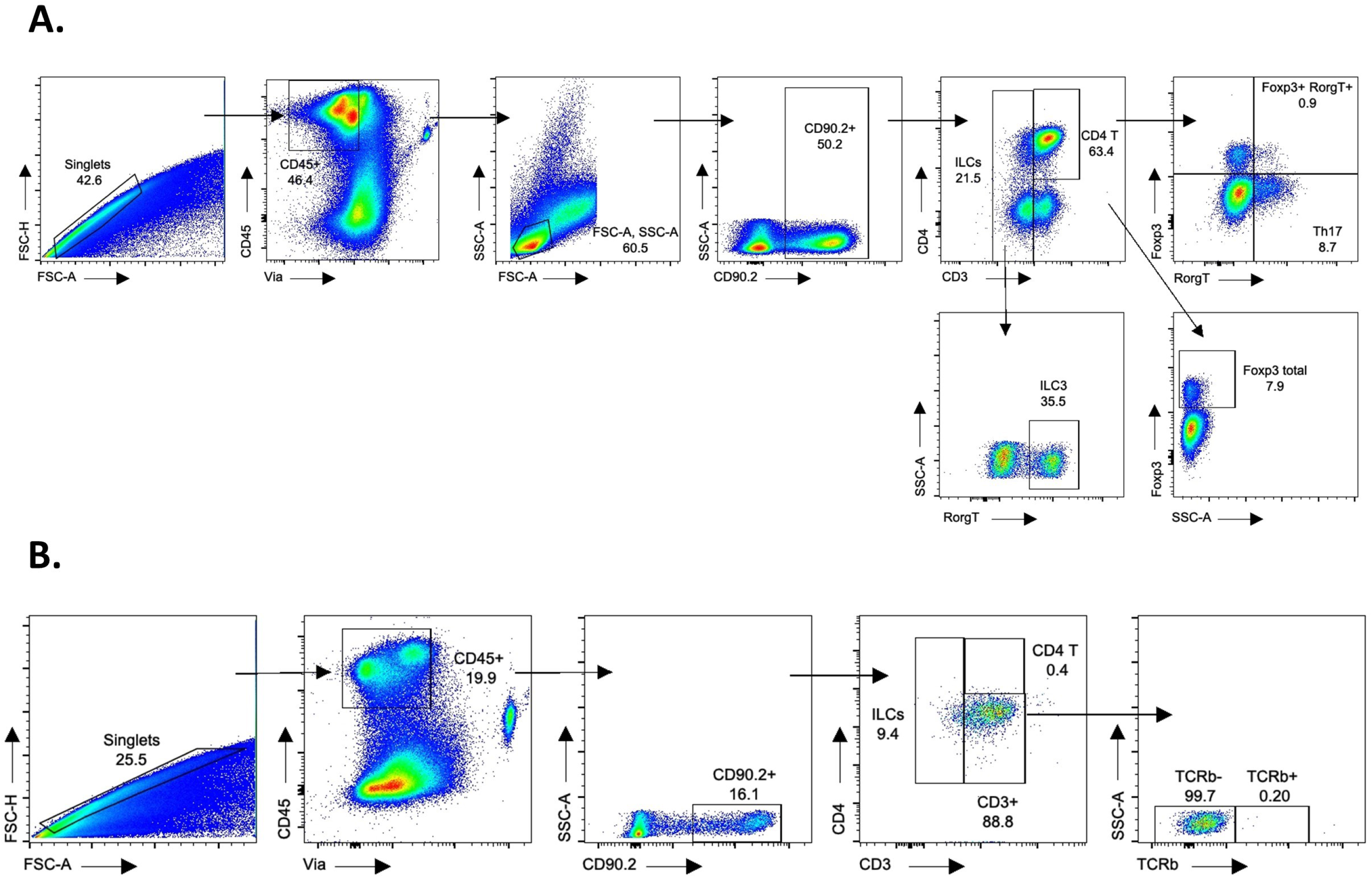
Flow cytometry gating strategies. **A**. The gating strategy that was employed to analyze ILCs, Th17 cells, total regulatory T cells and regulatory T cells expressing RORγT populations. Single and CD45^+^ live cells were only considered for the analysis. CD90.2^+^ lymphoid cells were gated with the forward scater/side scater (FSC/SSC) method. CD45^+^CD90.2^+^CD3^-^ cells were gated to analyze ILCs and the expression of RORγT served to distinguish ILC3 population. The CD45^+^CD90.2^+^CD3^+^ population was gated to analyze T cells and the Th17, total regulatory T cells and regulatory T cells expressing RORγT populations were analyzed based on the RORγT^+^ and FOXP3^+^ expression. **B.** The gating strategy that was employed to confirm T cell populations and ILC in T cell-deficient mice. Lamina propria cells from the ileum and colon were stained with CD45, viability, CD90, CD3, CD4, TCRβ. With this strategy, the first gate we isolated singlets to exclude doublets, following this, viable cells were gated using CD45 and viability. Then non-lymphocyte populations were deleted based on forward and side scater (FSC and SSC). Following this, CD90^+^ lymphoid cells were selected. The CD4^+^CD3^-^ show ILCs and CD4^+^CD3^+^TCRβ^+^gate distinguishes T cells.

**Supplementary Figure 6.**
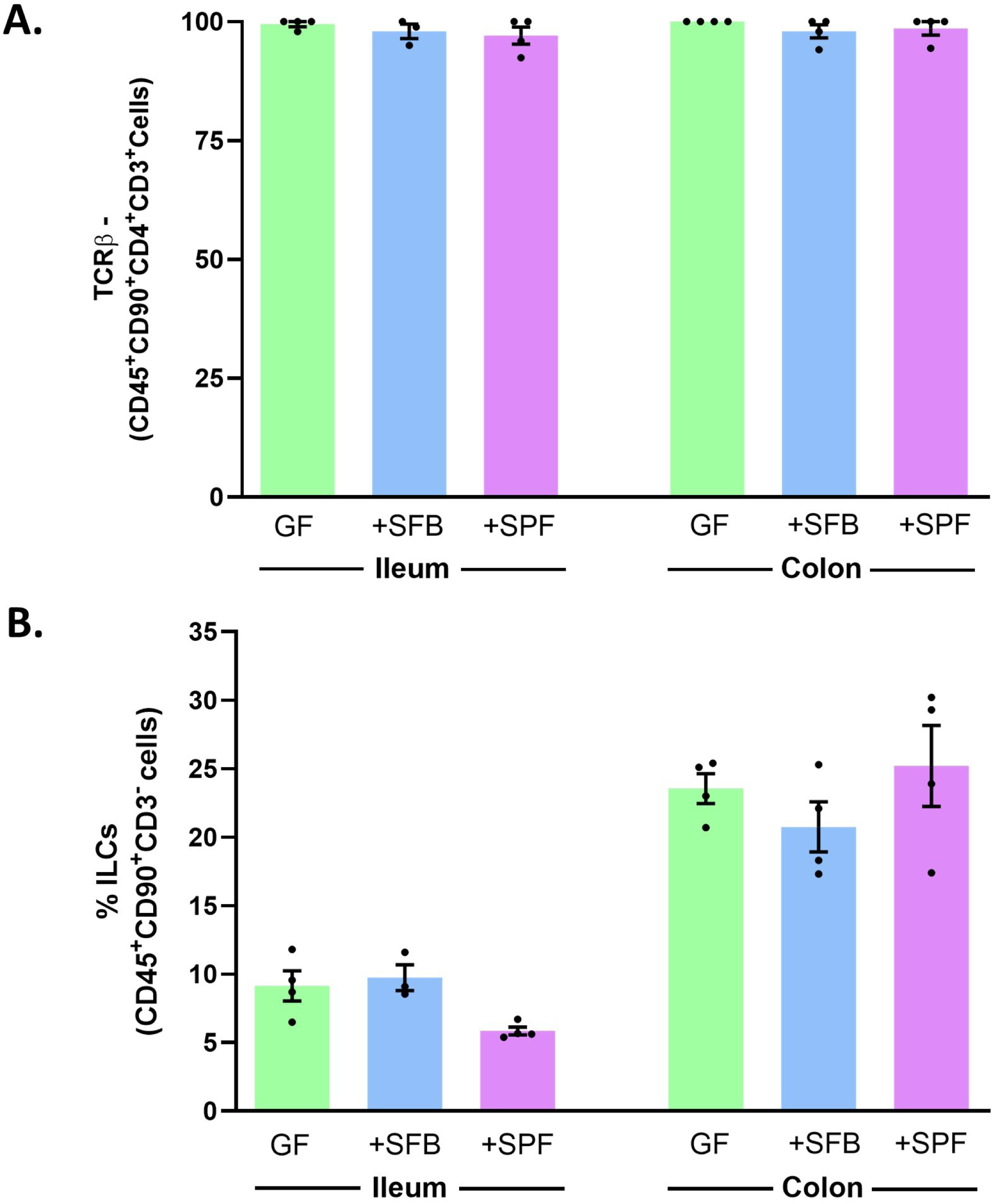
Confirmation of the phenotype of T cell-deficient mice. Colonization with segmented filamentous bacteria (SFB) in T cell deficient mice does not alter the expression of T cells and innate lymphoid cells (ILC). Germ-free male C57BL/6 and TCRβ^-^ deficient were colonized with SFB or with SPF microbiota and small intestinal lamina propria cells and colonic lamina propria cells were isolated. Graphs show the frequencies of TCRβ^-^ (A), and ILC (B) populations, all gated on CD45^+^CD90^+^ live cells. N=3-4 mice/group.

**Supplementary Figure 7.**
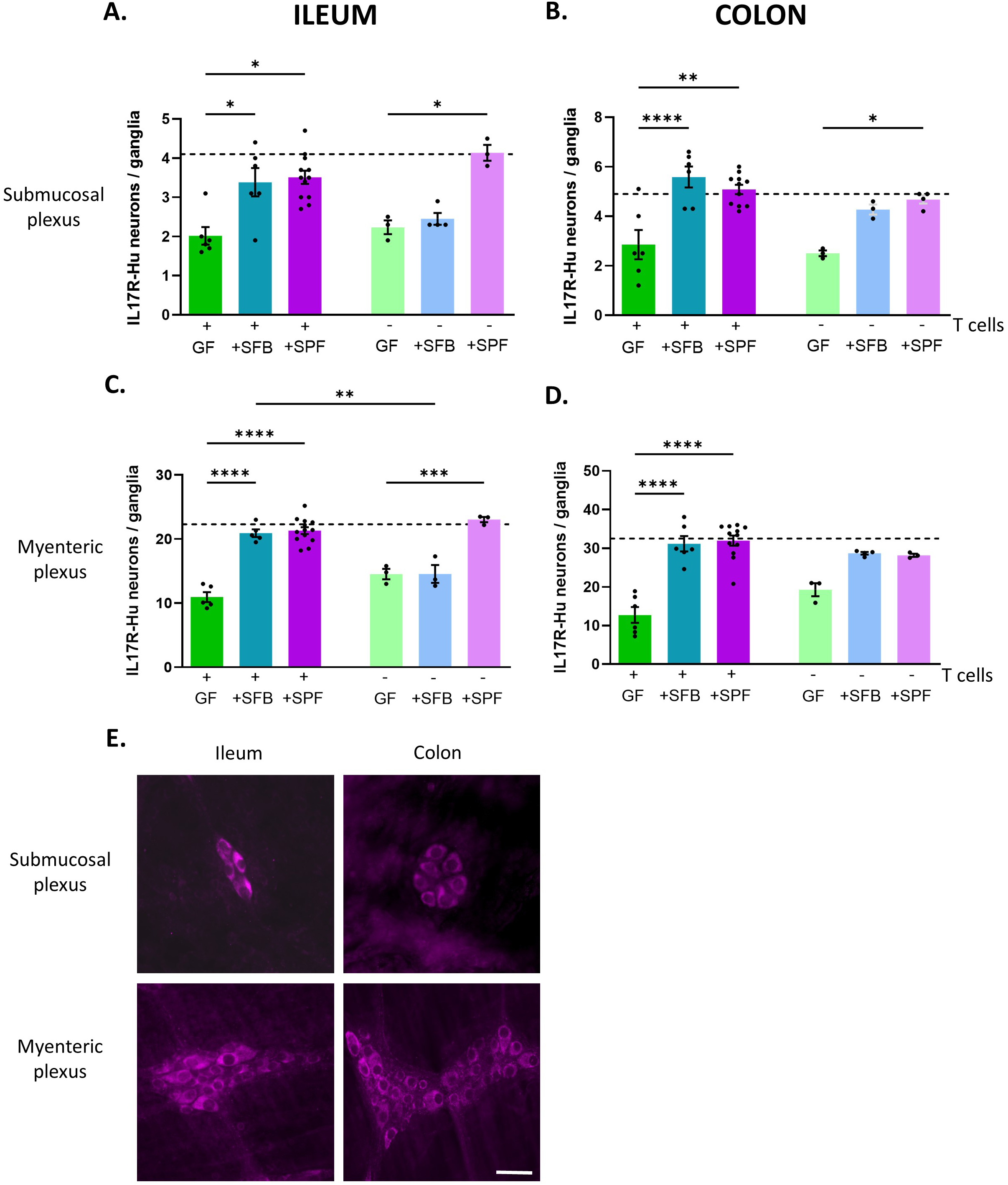
IL-17 receptor is expressed in enteric neurons. **A-D**. IL-17 receptor was localized using immunohistochemistry in neurons in the submucosal (**A, B**) and myenteric plexuses (**C, D**) in wildtype germ-free mice (GF), GF mice colonized with SFB or SPF microbiota (+), T cell-deficient GF mice, and T cell-deficient GF mice colonized with SFB or SPF microbiota (-) in the ileum (**A, C**) and colon (**B, D**). Each dot represents an individual animal. Dashed line indicates average neuronal density measured in wildtype mice colonized with SPF from birth. Data are pooled from 2-4 independent experiments run for each genotype. Data were analyzed using two-way ANOVA, followed by Šídák’s multiple comparison test. N=3-12/group. * p<0.05, ** p<0.01, *** p<0.001, **** p<0.0001. **E.** Representative immunohistochemical labeling of IL-17 receptor (purple) in the submucosal and myenteric plexus of the ileum and colon of wildtype germ-free mice colonized with SPF microbiota. Scale bar: 30μm.

**Supplementary Figure 8.**
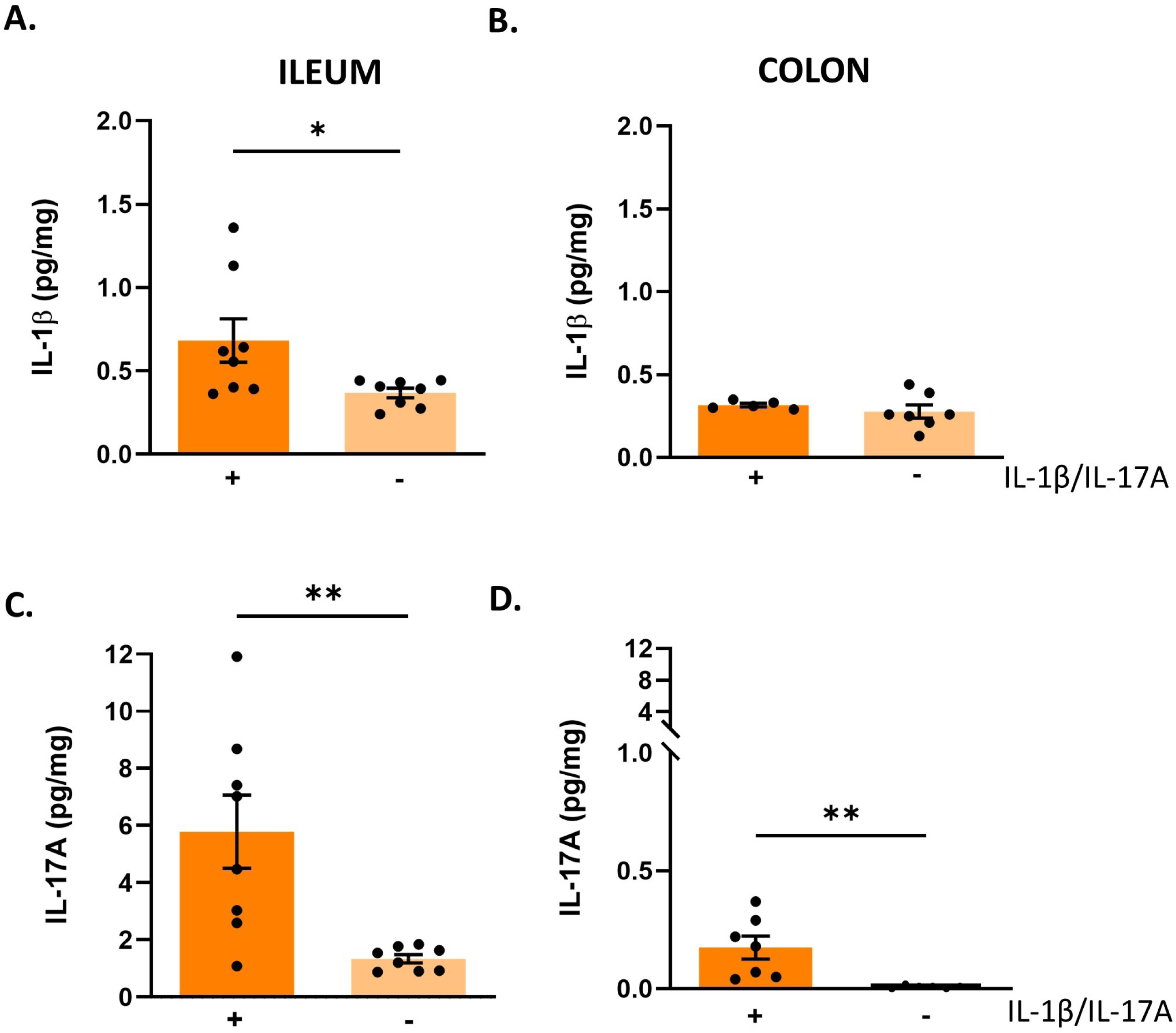
IL-1β and IL-17 levels are reduced by immunoneutralization. **A-B**. IL-1β levels in the ileum (**A**) and colon (**B**) of wildtype SPF colonized germ-free mice treated with isotype control antibodies (+) or a mixture of IL-1β and IL-17A neutralizing antibodies (-). **C-D.** IL-17A levels in the ileum (**C**) and colon (**C**) of wildtype SPF colonized germ-free mice treated with isotype control antibodies (+) or a mixture of IL-1β and IL-17A neutralizing antibodies (-). Each dot represents an individual animal. Data are pooled from 2 independent experiments. Data were analyzed with a Student’s t test. N=8/group. * p<0.05, ** p<0.01.

**Supplementary Figure 9.**
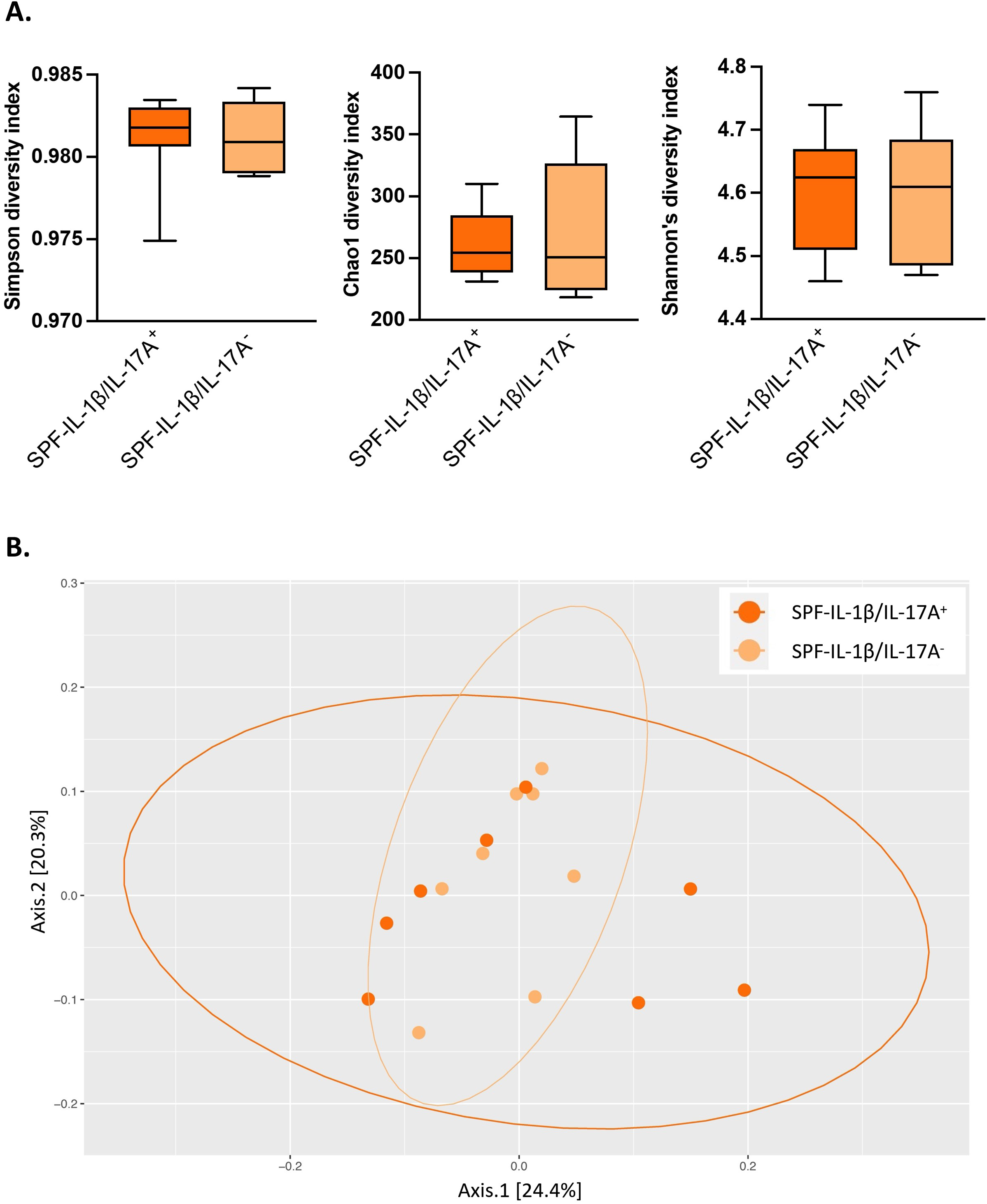
IL-1β and IL-17A immunoneutralization does not affect the microbiota composition of SPF colonized wildtype mice. **A**. Alpha Diversity was measured in samples wildtype germ-free mice colonized with SPF microbiota and treated with isotype control antibodies (SPF-IL-1β/IL-17A^+^) or neutralizing antibodies against IL-1β and IL-17A (SPF-IL-1β/IL-17A^-^) using the Simpson, Chao1, and Shannon Diversity Indices. Data are expressed as mean ± SEM, n= 8 mice per group, analyzed using either a Student’s t-test (Chao1, Shannon) or a Mann-Whitney test (Simpson) based on the distribution of the data. There were no statistical differences between the groups. **B.** Beta diversity was measured by generating a Bray-Curtis dissimilarity matrix, data was then plotted using a principal coordinate analysis plot. PERMANOVA statistical analysis was used for Bray-Curtis Dissimilarity to compare SPF-IL-1β/IL-17A^+^ and SPF-IL-1β/IL-17A^-^ groups, N=8 mice/group. Data are pooled from 2 independent experiments. There were no statistical differences between the groups.

**Supplementary Figure 10.**
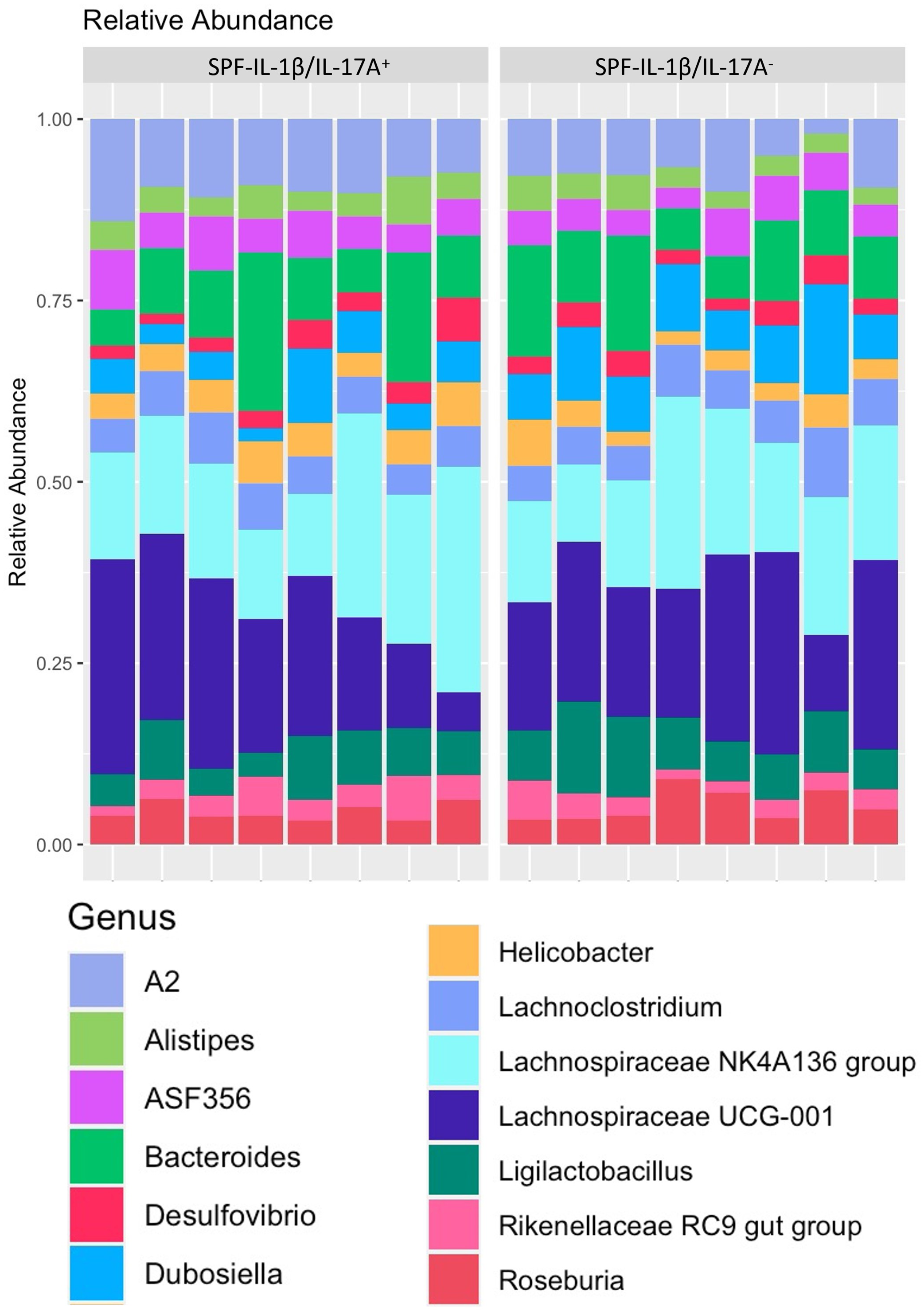
IL-1β and IL-17A immunoneutralization does not affect the top 30 prevalent amplicon sequencing variants (ASV). The top 30 prevalent ASVs were retrieved from wildtype germ-free mice colonized with SPF microbiota treated with isotype control antibodies (SPF-IL-1β/IL-17A^+^) or neutralizing antibodies against IL-1β and IL-17A (SPF-IL-1β/IL-17A^-^) and the relative abundance at the genus level was plotted per mouse.

**Supplementary Figure 11.**
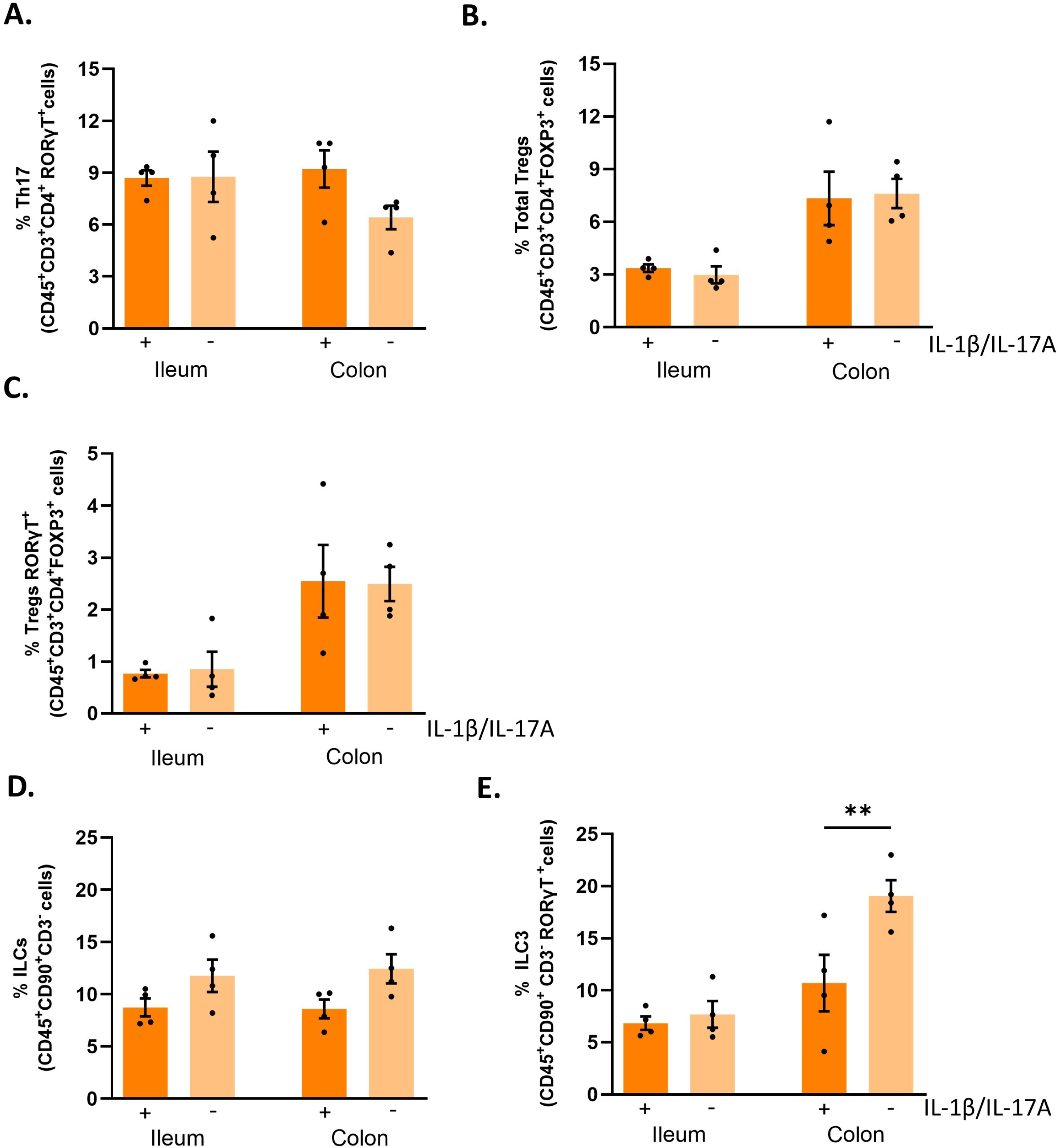
IL-1β and IL-17A immunoneutralization does not affect the populations of T cells in the lamina propria of the ileum and colon. Wildtype SPF colonized germ-free mice were treated with isotype control antibodies (+) or a mixture of IL-1β and IL-17A neutralizing antibodies (-) and small intestinal and large intestinal lamina propria cells were isolated. **A-C.** Frequencies of Th17 (**A**), total regulatory T cells (**B**) and regulatory T cells expressing RORγT (**C**) in the ileum and colon. Note that there were no significant differences between the groups. **D-F.** Frequencies of ILCs (**D**) and ILC3 cells (**E**) gated on CD45^+^CD90.2^+^CD3^-^. Note that there was a significant increase in frequency of ILC3 cells after treatment. Each dot represents an individual animal. Data are pooled from 2 independent experiments. Data were analyzed with a Student’s t test. N=4/group. ** p<0.01.

**Supplementary Figure 12.**
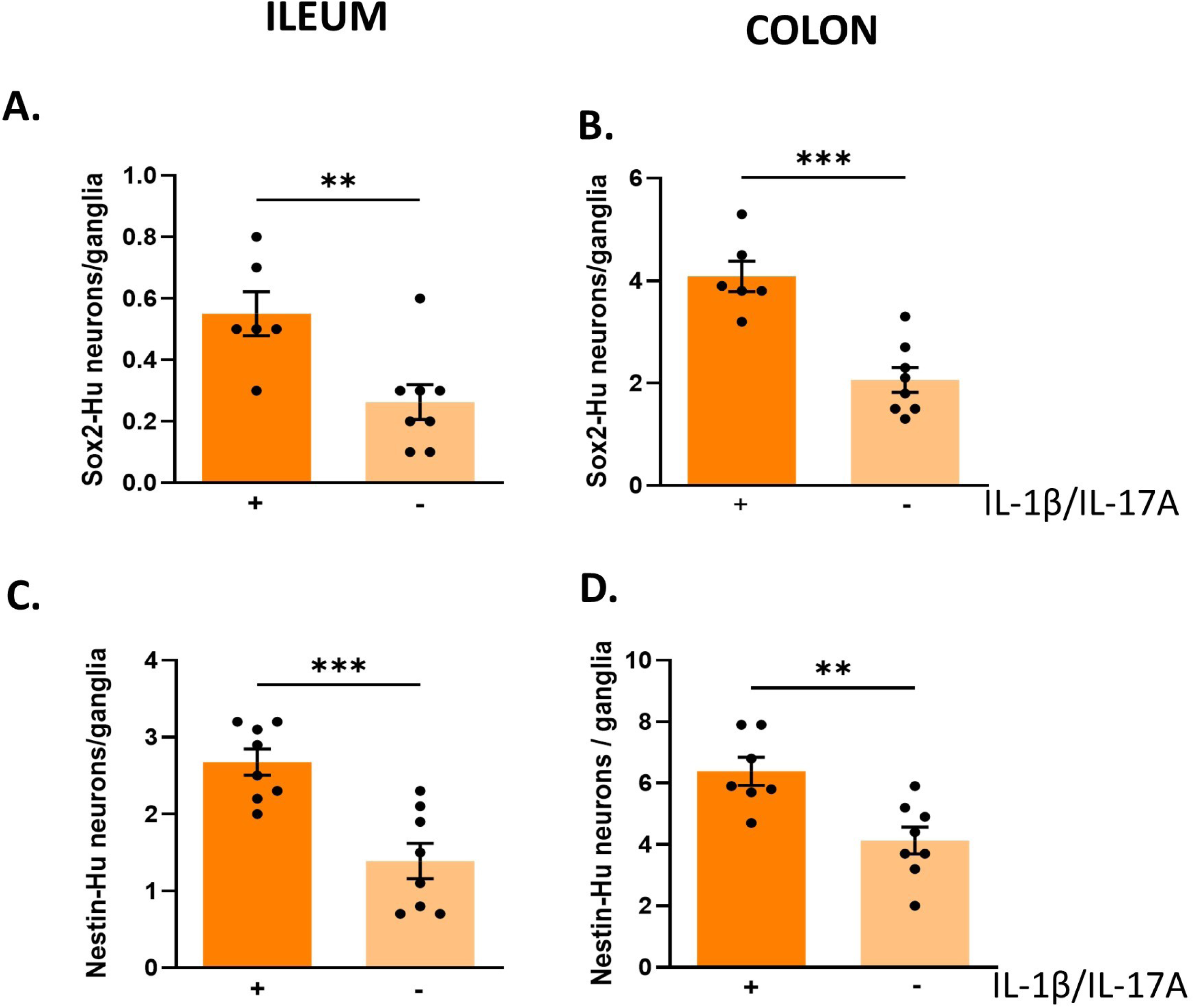
Sox2 and nestin are reduced in myenteric neurons after IL-1β and IL-17A immunoneutralization. **A-D**. Wildtype SPF colonized germ-free mice were treated with isotype control antibodies (+) or a mixture of IL-1β and IL-17A neutralizing antibodies (-). Sox2-HuC/D neurons in the myenteric plexus of the ileum (**A**) and colon (**B**) and nestin-HuC/D neurons in the myenteric plexus of the ileum (**C**) and colon (**D**). Each dot represents an individual animal. Data are pooled from 2 independent experiments. Data were analyzed with a Student’s t test. N=6-8/group. ** p<0.01, *** p<0.001.

**Supplementary Figure 13.**
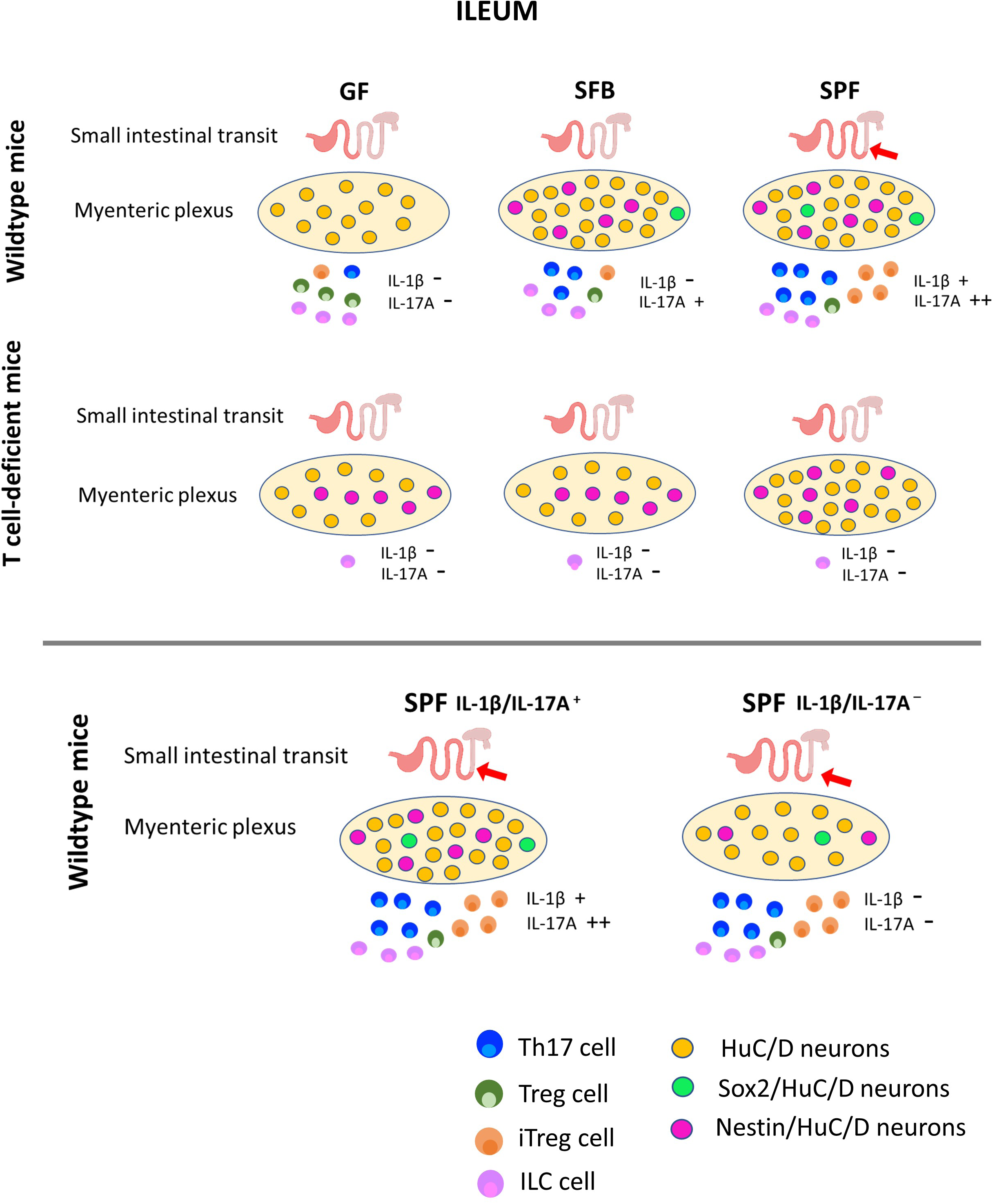
Graphic representation of the main findings in the ileum. **<u>Motility</u>**. Colonization with SPF normalized small intestinal motility in wildtype germ-free (GF) mice. SFB alone was not sufficient to restore normal motility in wildtype GF mice. T cells are necessary for the restoration of normal motility in GF mice colonized with SPF. Blocking IL-1β and IL-17A in GF mice colonized with SPF does not affect small intestinal motility. **<u>Enteric neurons</u>.** SFB and SPF microbiota are both sufficient to restore neuronal density in the ENS in wildtype GF mice. T cells are not required for the restoration of neuronal density by SPF in the ileum, but they are necessary for SFB to restore neuronal density. Both SFB and SFB stimulated Sox2 expression in the myenteric plexus of the ileum of wildtype mice. T cells are necessary for SFB and SPF stimulated Sox2 expression in the myenteric plexus of the ileum. Both SFB and SPF stimulated nestin expression in the myenteric plexus of the ileum of wildtype mice. The levels of nestin expression were higher in T cell-deficient mice, and there was no evidence that colonization altered the levels of expression in these mice. Immunoneutralization of IL-1β and IL-17A reduced neuronal density to levels similar to that of GF mice, and reduced Sox2 and nestin expression in SPF colonized mice. **<u>Immune cells and cytokines</u>.** Colonization with SFB and SPF markedly increased Th17 cells and Treg expressing RORγ+T cells in the ileum of wildtype GF mice. Colonization with SFB and SPF increased IL-1β and IL-17A in the ileum of wildtype GF mice, but not T cell-deficient mice. Immunoneutralization of IL-1β and IL-17A effectively reduced the levels of these cytokines without altering the populations of immune cells present in the lamina propria of the ileum.

**Supplementary Figure 14.**
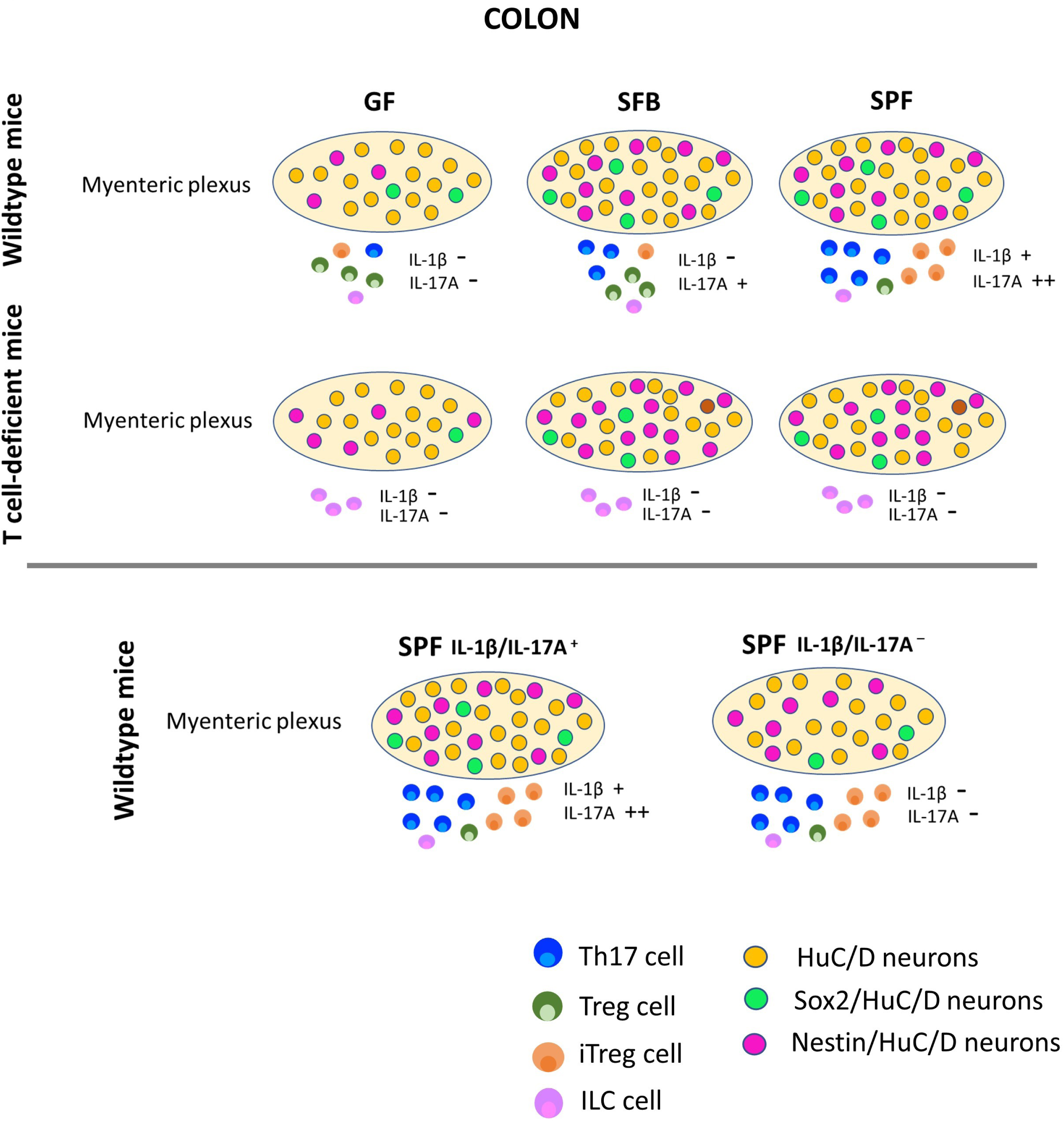
Graphic representation the main findings in the colon. **<u>Enteric neurons</u>.** SFB and SPF microbiota are both sufficient to restore neuronal density in the ENS in wildtype GF mice. T cells are not required for the restoration of neuronal density by either SFB or SPF. Both SFB and SPF stimulated Sox2 expression in the myenteric plexus of the colon of wildtype mice. T cells are not necessary for SFB and SPF stimulated Sox2 expression in the myenteric plexus of the colon. The levels of nestin expression were similar in all groups of mice and there was no evidence that colonization altered the levels of expression. Immunoneutralization of IL-1β and IL-17A reduced neuronal density, Sox2 and nestin expression in SPF colonized mice to levels similar to that of GF mice. **<u>Immune cells and cytokines</u>.** Colonization with SFB and SPF markedly increased Th17 cells and Treg expressing RORγ+T cells in the colon of wildtype GF mice. Colonization with SFB and SPF increased IL-1β and IL-17A in the colon of wildtype GF mice, but not T cell-deficient mice. Immunoneutralization of IL-1β and IL-17A effectively reduced the levels of these cytokines without altering the populations of immune cells present in the lamina propria of the colon.

## Supplementary Results

### Body weight

Wildtype germ-free mice (aged 10-13 weeks) colonized with either SFB or SPF had significantly reduced body weight (26.7±1.0g vs 22.7±0.5g and 23.2±0.4g respectively; p<0.001 GF vs SFB and p<0.01 GF vs SPF; two-way ANOVA [n=10/group]). In the T-cell deficient germ-free mice (aged 13-23 weeks), colonization with SFB did not result in a reduction in body weight compared to germ-free mice, as was seen in wild-type SFB-colonized germ-free mice. However, the T cell-deficient SPF-colonized mice had a significantly reduced body weight compared to both the germ-free and SFB monocolonized mice, similar to what was observed in wild-type SPF-colonized mice (22.3±0.5g vs 25.9±0.5 and 26.8±0.7g respectively, p<0.01 GF vs SPF and p<0.001 SFB vs SPF, two-way ANOVA, [n=8/group]). There was a significant difference in weight between SFB moncolonized wildtype germ-free mice compared to the T-cell deficient germ-free mice (p<0.001, two-way ANOVA).

Treatment with IL-1β/IL-17A neutralizing antibodies had nο effect on body weight (23.8 ± 0.6g isotype controls, 24.4 ± 0.5g IL-1β/IL-17A treated mice [n=8]).

### Colon length

The colon length in the SPF colonized mice was similar to that of germ-free or SFB monocolonized mice (7.8 ± 0.2cm vs 8.6 ± 0.2cm and 8.4 ± 0.3cm, respectively; p>0.05, two-way ANOVA, [n=10/group]). The length of the colon of the SPF-colonized mice is the same as that of mice raised under SPF conditions from birth (7.8 ± 0.2cm vs 7.7 ± 0.1cm [n=10]). There was a small, but significantly reduced colon length in the T cell-deficient SPF colonized mice compared to germ-free T-cell deficient mice (7.3±0.3cm, vs 8.4±0.3cm, [n=8/group], p<0.05, two-way ANOVA). SFB colonized T cell-deficient mice had a colon length identical to germ-free T-cell deficient mice (8.4±0.4cm, [n=8/group]). There were no significant differences in colon length between wildtype germ-free mice compared to the T-cell deficient germ-free mice (p>0.05, two-way ANOVA).

### Intestinal cytokines

The levels of cytokines in the wall of the ileum and colon are found in Supplementary Tables 1 and Table 2. There was an inverse correlation of lipopolysaccharide-induced CXC chemokine 5 (LIX) levels and Sox2-HuC/D in the myenteric plexus of the ileum and colon of wild-type mice, but this only extended to the levels observed in the colon of T cell-deficient mice. The cytokines, IFNγ, IL-1α, MCP-1 and MIP-2 were elevated in both the ileum and colon of T cell-deficient mice and these correlate with elevated nestin expression in HuC/D neurons in the myenteric plexus of the ileum and colon in these mice.

We also noted some other specific patterns of cytokine expression correlated with the colonization of the mice. In wild-type mice, in the ileum, the levels of IL-17A, and LIX (inversely) correlated with presence of SFB and SPF microbiota. In T cell-deficient mice, in the ileum, we observed elevated levels of GM-CSF and TNF that corresponded to the microbial colonization and a negative correlation with IL-12p70 levels. In the colon in wild-type mice, we did not observe any positive correlation of cytokines with both SFB and SPF microbiota, but we saw a negative correlation with IL-10 and LIX levels. However, we saw that levels of GM-CSF, IFNγ, IL-1β, IL-17A and TNFα correlated with the presence of SPF microbiota. In the colon of T cell-deficient mice, we did not observe any correlations of cytokines with both SFB and SPF microbiota. However, we saw a positive correlation of the levels of GM-CSF, IL-1β, IL-6, KC, MCP-1 and TNF with SPF microbiota and a negative correlation with the levels of IL-2, IL-4, IL-7, IL-10, IL-13 and LIX.

## Supplementary Tables

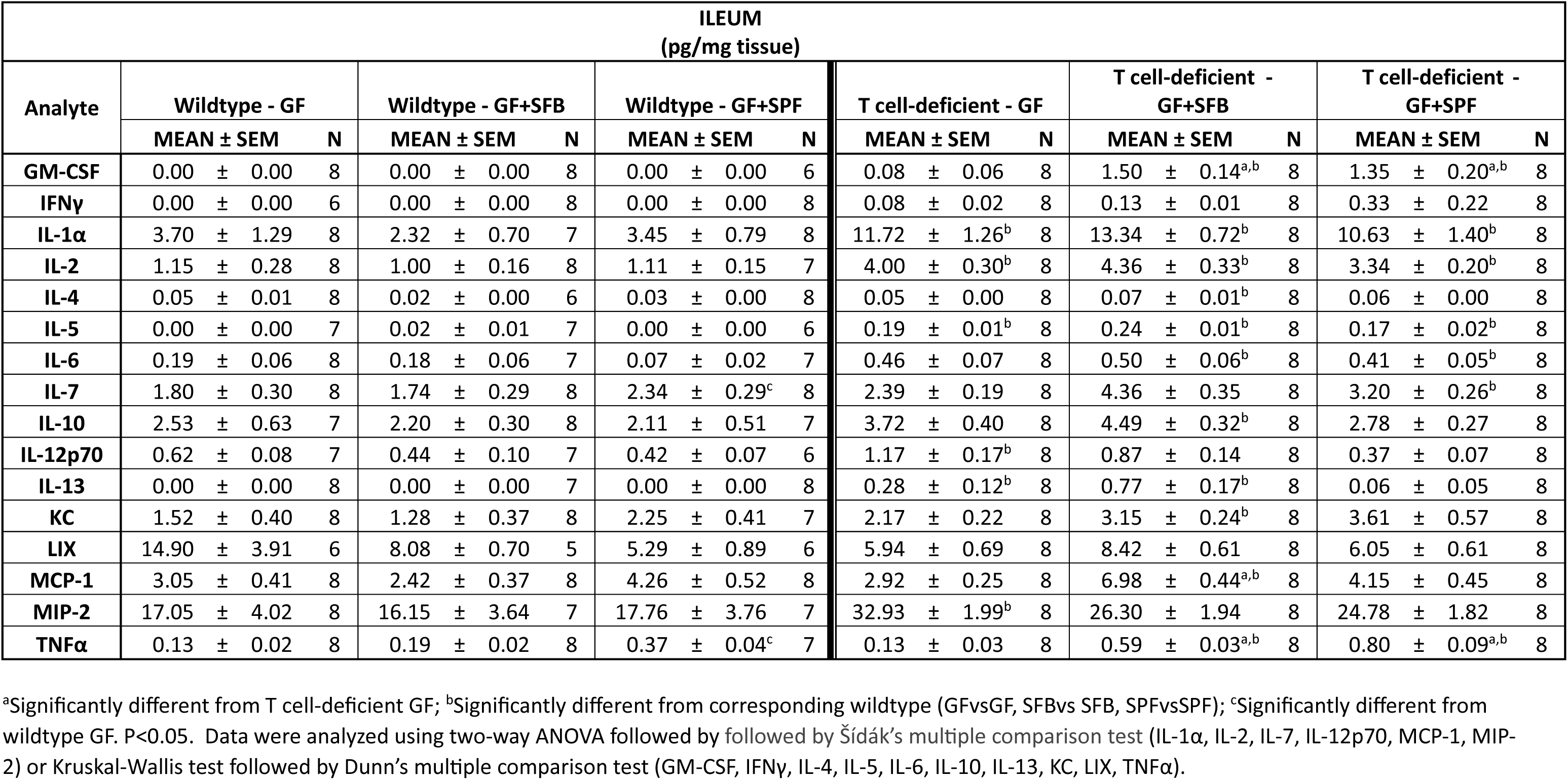
Supplementary Table 1.

Supplementary Table 1
| ILEUM<br>(pg/mg tissue) |  |  |  |  |  |  |  |  |  |  |  |  |  |  |  |  |  |  |
| --- | --- | --- | --- | --- | --- | --- | --- | --- | --- | --- | --- | --- | --- | --- | --- | --- | --- | --- |
| Analyte | Wildtype - GF |  |  | Wildtype - GF+SFB |  |  | Wildtype - GF+SPF |  |  | T cell-deficient - GF |  |  | T cell-deficient - GF+SFB |  |  | T cell-deficient - GF+SPF |  |  |
|  | MEAN ± SEM |  | N | MEAN ± SEM |  | N | MEAN ± SEM |  | N | MEAN ± SEM |  | N | MEAN ± SEM |  | N | MEAN ± SEM |  | N |
| GM-CSF | 0.00 | ± 0.00 | 8 | 0.00 | ± 0.00 | 8 | 0.00 | ± 0.00 | 6 | 0.08 | ± 0.06 | 8 | 1.50 | ± 0.14<sup>a,b</sup> | 8 | 1.35 | ± 0.20<sup>a,b</sup> | 8 |
| IFNγ | 0.00 | ± 0.00 | 6 | 0.00 | ± 0.00 | 8 | 0.00 | ± 0.00 | 8 | 0.08 | ± 0.02 | 8 | 0.13 | ± 0.01 | 8 | 0.33 | ± 0.22 | 8 |
| IL-1α | 3.70 | ± 1.29 | 8 | 2.32 | ± 0.70 | 7 | 3.45 | ± 0.79 | 8 | 11.72 | ± 1.26<sup>b</sup> | 8 | 13.34 | ± 0.72<sup>b</sup> | 8 | 10.63 | ± 1.40<sup>b</sup> | 8 |
| IL-2 | 1.15 | ± 0.28 | 8 | 1.00 | ± 0.16 | 8 | 1.11 | ± 0.15 | 7 | 4.00 | ± 0.30<sup>b</sup> | 8 | 4.36 | ± 0.33<sup>b</sup> | 8 | 3.34 | ± 0.20<sup>b</sup> | 8 |
| IL-4 | 0.05 | ± 0.01 | 8 | 0.02 | ± 0.00 | 6 | 0.03 | ± 0.00 | 8 | 0.05 | ± 0.00 | 8 | 0.07 | ± 0.01<sup>b</sup> | 8 | 0.06 | ± 0.00 | 8 |
| IL-5 | 0.00 | ± 0.00 | 7 | 0.02 | ± 0.01 | 7 | 0.00 | ± 0.00 | 6 | 0.19 | ± 0.01<sup>b</sup> | 8 | 0.24 | ± 0.01<sup>b</sup> | 8 | 0.17 | ± 0.02<sup>b</sup> | 8 |
| IL-6 | 0.19 | ± 0.06 | 8 | 0.18 | ± 0.06 | 7 | 0.07 | ± 0.02 | 7 | 0.46 | ± 0.07 | 8 | 0.50 | ± 0.06<sup>b</sup> | 8 | 0.41 | ± 0.05<sup>b</sup> | 8 |
| IL-7 | 1.80 | ± 0.30 | 8 | 1.74 | ± 0.29 | 8 | 2.34 | ± 0.29<sup>c</sup> | 8 | 2.39 | ± 0.19 | 8 | 4.36 | ± 0.35 | 8 | 3.20 | ± 0.26<sup>b</sup> | 8 |
| IL-10 | 2.53 | ± 0.63 | 7 | 2.20 | ± 0.30 | 8 | 2.11 | ± 0.51 | 7 | 3.72 | ± 0.40 | 8 | 4.49 | ± 0.32<sup>b</sup> | 8 | 2.78 | ± 0.27 | 8 |
| IL-12p70 | 0.62 | ± 0.08 | 7 | 0.44 | ± 0.10 | 7 | 0.42 | ± 0.07 | 6 | 1.17 | ± 0.17<sup>b</sup> | 8 | 0.87 | ± 0.14 | 8 | 0.37 | ± 0.07 | 8 |
| IL-13 | 0.00 | ± 0.00 | 8 | 0.00 | ± 0.00 | 7 | 0.00 | ± 0.00 | 8 | 0.28 | ± 0.12<sup>b</sup> | 8 | 0.77 | ± 0.17<sup>b</sup> | 8 | 0.06 | ± 0.05 | 8 |
| KC | 1.52 | ± 0.40 | 8 | 1.28 | ± 0.37 | 8 | 2.25 | ± 0.41 | 7 | 2.17 | ± 0.22 | 8 | 3.15 | ± 0.24<sup>b</sup> | 8 | 3.61 | ± 0.57 | 8 |
| LIX | 14.90 | ± 3.91 | 6 | 8.08 | ± 0.70 | 5 | 5.29 | ± 0.89 | 6 | 5.94 | ± 0.69 | 8 | 8.42 | ± 0.61 | 8 | 6.05 | ± 0.61 | 8 |
| MCP-1 | 3.05 | ± 0.41 | 8 | 2.42 | ± 0.37 | 8 | 4.26 | ± 0.52 | 8 | 2.92 | ± 0.25 | 8 | 6.98 | ± 0.44<sup>a,b</sup> | 8 | 4.15 | ± 0.45 | 8 |
| MIP-2 | 17.05 | ± 4.02 | 8 | 16.15 | ± 3.64 | 7 | 17.76 | ± 3.76 | 7 | 32.93 | ± 1.99<sup>b</sup> | 8 | 26.30 | ± 1.94 | 8 | 24.78 | ± 1.82 | 8 |
| TNFα | 0.13 | ± 0.02 | 8 | 0.19 | ± 0.02 | 8 | 0.37 | ± 0.04<sup>c</sup> | 7 | 0.13 | ± 0.03 | 8 | 0.59 | ± 0.03<sup>a,b</sup> | 8 | 0.80 | ± 0.09<sup>a,b</sup> | 8 |

<sup>a</sup>Significantly different from T cell-deficient GF; <sup>b</sup>Significantly different from corresponding wildtype (GFvsGF, SFBvs SFB, SPFvsSPF); <sup>c</sup>Significantly different from wildtype GF. P<0.05. Data were analyzed using two-way ANOVA followed by followed by Šídák's multiple comparison test (IL-1α , IL-2, IL-7, IL-12p70, MCP-1, MIP-2) or Kruskal-Wallis test followed by Dunn's multiple comparison test (GM-CSF, IFNγ , IL-4, IL-5, IL-6, IL-10, IL-13, KC, LIX, TNFα ).

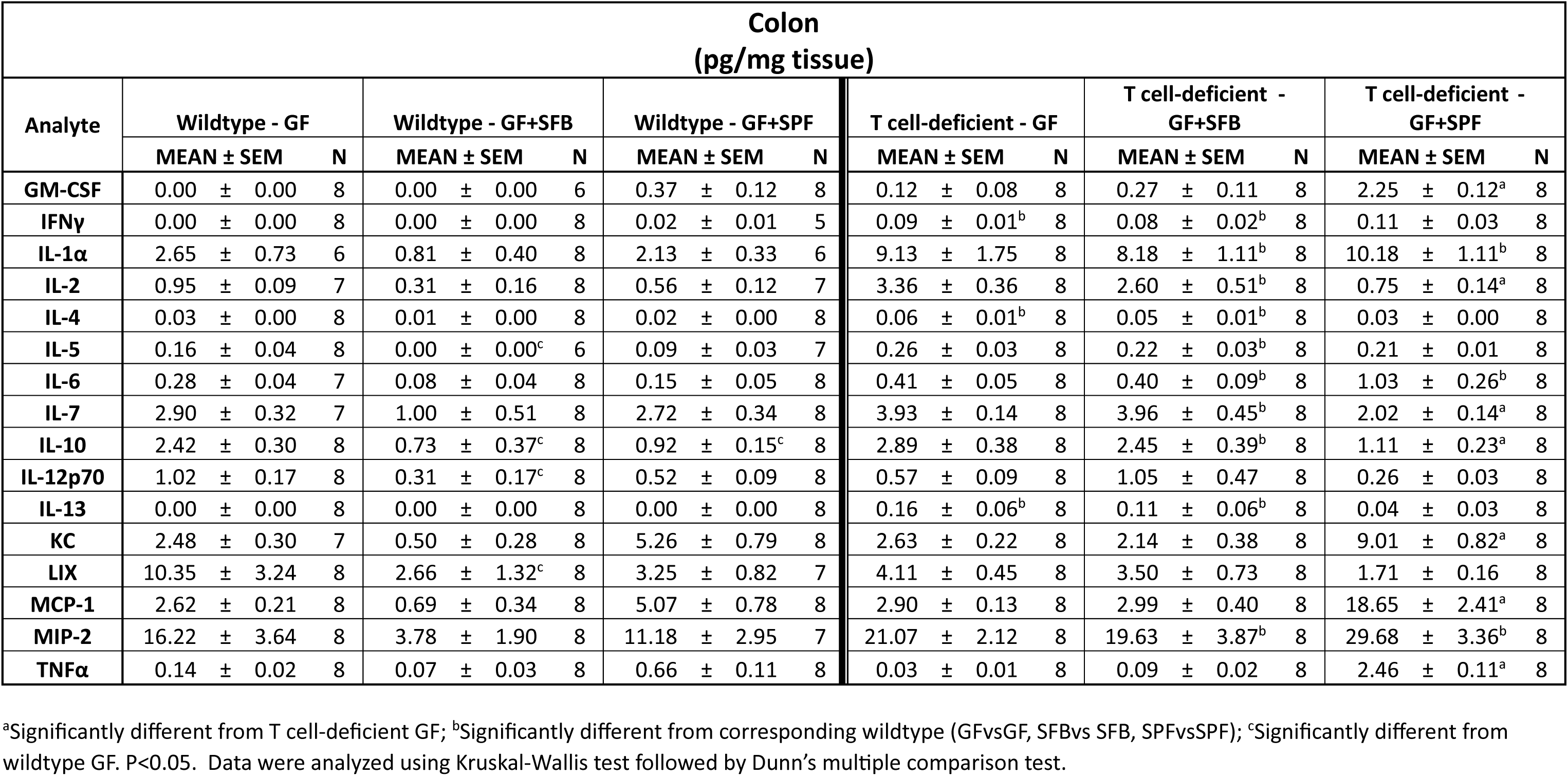
Supplementary Table 2.

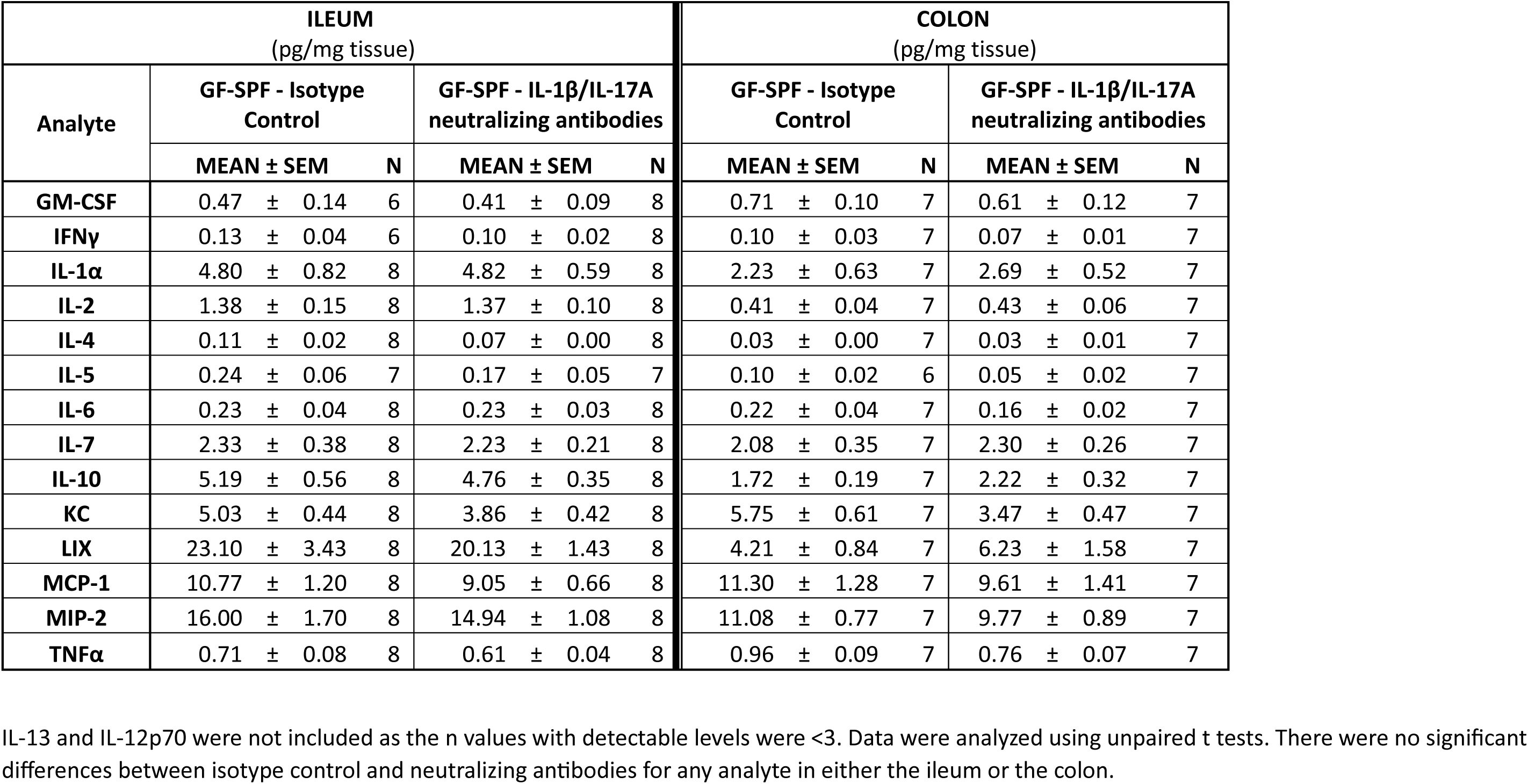
Supplementary Table 3.

## References

1. Sharkey, K. A., Mawe, G. M. The enteric nervous system. Physiol Rev 103, 1487–1564 (2023).

2. Furness, J. B. The enteric nervous system and neurogastroenterology. Nat Rev Gastroenterol Hepatol 9, 286–294 (2012).

3. Seguella, L., Gulbransen, B. D. Enteric glial biology, intercellular signalling and roles in gastrointestinal disease. Nat Rev Gastroenterol Hepatol 18, 571–587 (2021).

4. Joly, A., Leulier, F., De Vadder, F. Microbial Modulation of the Development and Physiology of the Enteric Nervous System. Trends Microbiol 29, 686–699 (2021).

5. Gershon, M. D., Margolis, K. G. The gut, its microbiome, and the brain: connections and communications. J Clin Invest 131, (2021).

6. Cryan, J. F., et al. The Microbiota-Gut-Brain Axis. Physiol Rev 99, 1877–2013 (2019).

7. Griffiths, J. A., et al. Peripheral neuronal activation shapes the microbiome and alters gut physiology. Cell Rep 43, 113953 (2024).

8. Vicentini, F. A., et al. Intestinal microbiota shapes gut physiology and regulates enteric neurons and glia. Microbiome 9, 210 (2021).

9. Obata, Y., et al. Neuronal programming by microbiota regulates intestinal physiology. Nature 578, 284–289 (2020).

10. Bernabe, G., et al. Antibiotic Treatment Induces Long-Lasting Effects on Gut Microbiota and the Enteric Nervous System in Mice. Antibiotics (Basel) 12, (2023).

11. Bai, X., et al. Vasoactive Intestinal Polypeptide Plays a Key Role in the Microbial-Neuroimmune Control of Intestinal Motility. Cell Mol Gastroenterol Hepatol 17, 383–398 (2024).

12. Caputi, V., et al. Antibiotic-induced dysbiosis of the microbiota impairs gut neuromuscular function in juvenile mice. Br J Pharmacol 174, 3623–3639 (2017).

13. Muller, P. A., et al. Crosstalk between muscularis macrophages and enteric neurons regulates gastrointestinal motility. Cell 158, 300–313 (2014).

14. Muller, P. A., Matheis, F., Schneeberger, M., Kerner, Z., Jove, V., Mucida, D. Microbiota-modulated CART(+) enteric neurons autonomously regulate blood glucose. Science 370, 314–321 (2020).

15. De Vadder, F., et al. Gut microbiota regulates maturation of the adult enteric nervous system via enteric serotonin networks. Proc Natl Acad Sci U S A 115, 6458–6463 (2018).

16. Kashyap, P. C., et al. Complex interactions among diet, gastrointestinal transit, and gut microbiota in humanized mice. Gastroenterology 144, 967–977 (2013).

17. Frith, M. E., Kashyap, P. C., Linden, D. R., Theriault, B., Chang, E. B. Microbiota-dependent early life programming of gastrointestinal motility. bioRxiv, (2023).

18. Yano, J. M., et al. Indigenous bacteria from the gut microbiota regulate host serotonin biosynthesis. Cell 161, 264–276 (2015).

19. Yan, Y., et al. Interleukin-6 produced by enteric neurons regulates the number and phenotype of microbe-responsive regulatory T cells in the gut. Immunity 54, 499–513 e495 (2021).

20. Collins, J., Borojevic, R., Verdu, E. F., Huizinga, J. D., Ratcliffe, E. M. Intestinal microbiota influence the early postnatal development of the enteric nervous system. Neurogastroenterol Motil 26, 98–107 (2014).

21. Ignacio, A., et al. Small intestinal resident eosinophils maintain gut homeostasis following microbial colonization. Immunity 55, 1250–1267 e1212 (2022).

22. Jarret, A., et al. Enteric Nervous System-Derived IL-18 Orchestrates Mucosal Barrier Immunity. Cell 180, 50–63 e12 (2020).

23. Rahman, A. A., et al. Optogenetic Activation of Cholinergic Enteric Neurons Reduces Inflammation in Experimental Colitis. Cell Mol Gastroenterol Hepatol 17, 907–921 (2024).

24. Balasubramaniam, A., Srinivasan, S. Role of stimulator of interferon genes (STING) in the enteric nervous system in health and disease. Neurogastroenterol Motil 35, e14603 (2023).

25. Drokhlyansky, E., et al. The Human and Mouse Enteric Nervous System at Single-Cell Resolution. Cell 182, 1606–1622 e1623 (2020).

26. Zeisel, A., et al. Molecular Architecture of the Mouse Nervous System. Cell 174, 999–1014 e1022 (2018).

27. Kulkarni, S., et al. Adult enteric nervous system in health is maintained by a dynamic balance between neuronal apoptosis and neurogenesis. Proc Natl Acad Sci U S A 114, E3709–E3718 (2017).

28. Belkind-Gerson, J., et al. Colitis induces enteric neurogenesis through a 5-HT4-dependent mechanism. Inflamm Bowel Dis 21, 870–878 (2015).

29. Laranjeira, C., et al. Glial cells in the mouse enteric nervous system can undergo neurogenesis in response to injury. J Clin Invest 121, 3412–3424 (2011).

30. Belkind-Gerson, J., et al. Colitis promotes neuronal differentiation of Sox2+ and PLP1+ enteric cells. Sci Rep 7, 2525 (2017).

31. Grundmann, D., Markwart, F., Scheller, A., Kirchhoff, F., Schafer, K. H. Phenotype and distribution pattern of nestin-GFP-expressing cells in murine myenteric plexus. Cell Tissue Res 366, 573–586 (2016).

32. Joseph, N. M., He, S., Quintana, E., Kim, Y. G., Nunez, G., Morrison, S. J. Enteric glia are multipotent in culture but primarily form glia in the adult rodent gut. J Clin Invest 121, 3398–3411 (2011).

33. Middelhoff, M., et al. Adult enteric Dclk1-positive glial and neuronal cells reveal distinct responses to acute intestinal injury. Am J Physiol Gastrointest Liver Physiol 322, G583–G597 (2022).

34. Yarandi, S. S., Kulkarni, S., Saha, M., Sylvia, K. E., Sears, C. L., Pasricha, P. J. Intestinal Bacteria Maintain Adult Enteric Nervous System and Nitrergic Neurons via Toll-like Receptor 2-induced Neurogenesis in Mice. Gastroenterology 159, 200–213 e208 (2020).

35. Cantarero Carmona, I., Luesma Bartolome, M. J., Lavoie-Gagnon, C., Junquera Escribano, C. Distribution of nestin protein: immunohistochemical study in enteric plexus of rat duodenum. Microsc Res Tech 74, 148–152 (2011).

36. Ivanov, II, et al. Induction of intestinal Th17 cells by segmented filamentous bacteria. Cell 139, 485–498 (2009).

37. Davis, C. P., Savage, D. C. Habitat, succession, attachment, and morphology of segmented, filamentous microbes indigenous to the murine gastrointestinal tract. Infect Immun 10, 948–956 (1974).

38. Kosiewicz, M. M., Zirnheld, A. L., Alard, P. Gut microbiota, immunity, and disease: a complex relationship. Front Microbiol 2, 180 (2011).

39. Han, L., et al. Innate Lymphoid Cells: A Link between the Nervous System and Microbiota in Intestinal Networks. Mediators Inflamm 2019, 1978094 (2019).

40. Jakob, M. O., Murugan, S., Klose, C. S. N. Neuro-Immune Circuits Regulate Immune Responses in Tissues and Organ Homeostasis. Front Immunol 11, 308 (2020).

41. Ghaedi, M., Takei, F. Innate lymphoid cell development. J Allergy Clin Immunol 147, 1549–1560 (2021).

42. Cherrier, D. E., Serafini, N., Di Santo, J. P. Innate Lymphoid Cell Development: A T Cell Perspective. Immunity 48, 1091–1103 (2018).

43. Tan, T. G., et al. Identifying species of symbiont bacteria from the human gut that, alone, can induce intestinal Th17 cells in mice. Proc Natl Acad Sci U S A 113, E8141–E8150 (2016).

44. Hooper, L. V., Littman, D. R., Macpherson, A. J. Interactions between the microbiota and the immune system. Science 336, 1268–1273 (2012).

45. Maynard, C. L., Elson, C. O., Hatton, R. D., Weaver, C. T. Reciprocal interactions of the intestinal microbiota and immune system. Nature 489, 231–241 (2012).

46. Poon, S. S. B., et al. Neonatal antibiotics have long term sex-dependent effects on the enteric nervous system. J Physiol 600, 4303–4323 (2022).

47. McVey Neufeld, K. A., Perez-Burgos, A., Mao, Y. K., Bienenstock, J., Kunze, W. A. The gut microbiome restores intrinsic and extrinsic nerve function in germ-free mice accompanied by changes in calbindin. Neurogastroenterol Motil 27, 627–636 (2015).

48. Anitha, M., Vijay-Kumar, M., Sitaraman, S. V., Gewirtz, A. T., Srinivasan, S. Gut microbial products regulate murine gastrointestinal motility via Toll-like receptor 4 signaling. Gastroenterology 143, 1006–1016 e1004 (2012).

49. Hung, L. Y., et al. Neonatal Antibiotics Disrupt Motility and Enteric Neural Circuits in Mouse Colon. Cell Mol Gastroenterol Hepatol 8, 298–300 e296 (2019).

50. Aktar, R., et al. Human resident gut microbe Bacteroides thetaiotaomicron regulates colonic neuronal innervation and neurogenic function. Gut Microbes 11, 1745–1757 (2020).

51. Belkaid, Y., Harrison, O. J. Homeostatic Immunity and the Microbiota. Immunity 46, 562–576 (2017).

52. Ansaldo, E., Farley, T. K., Belkaid, Y. Control of Immunity by the Microbiota. Annu Rev Immunol 39, 449–479 (2021).

53. Geuking, M. B., et al. Intestinal bacterial colonization induces mutualistic regulatory T cell responses. Immunity 34, 794–806 (2011).

54. Mori, D., et al. IL-17A induces hypo-contraction of intestinal smooth muscle via induction of iNOS in muscularis macrophages. J Pharmacol Sci 125, 394–405 (2014).

55. Li, J., et al. Targeting IL-17A Improves the Dysmotility of the Small Intestine and Alleviates the Injury of the Interstitial Cells of Cajal during Sepsis. Oxid Med Cell Longev 2019, 1475729 (2019).

56. Vallance, B. A., Collins, S. M. The effect of nematode infection upon intestinal smooth muscle function. Parasite Immunol 20, 249–253 (1998).

57. Vicentini, F. A., Fahlman, T., Raptis, S. G., Wallace, L. E., Hirota, S. A., Sharkey, K. A. New Concepts of the Interplay Between the Gut Microbiota and the Enteric Nervous System in the Control of Motility. Adv Exp Med Biol 1383, 55–69 (2022).

58. McKay, D. M., Defaye, M., Rajeev, S., MacNaughton, W. K., Nasser, Y., Sharkey, K. A. Neuroimmunophysiology of the Gastrointestinal Tract. Am J Physiol Gastrointest Liver Physiol, (2024).

59. Mombaerts, P., et al. Mutations in T-cell antigen receptor genes alpha and beta block thymocyte development at different stages. Nature 360, 225–231 (1992).

60. Prakash, T., et al. Complete genome sequences of rat and mouse segmented filamentous bacteria, a potent inducer of th17 cell differentiation. Cell Host Microbe 10, 273–284 (2011).

61. Quast, C., et al. The SILVA ribosomal RNA gene database project: improved data processing and web-based tools. Nucleic Acids Res 41, D590–596 (2013).

